# Whole-proteome structure/function prediction in Uropathogenic *E. coli* reveals previously missed host-microbe and microbe-phage interaction pathways

**DOI:** 10.64898/2026.06.16.732597

**Authors:** Chunxiang Peng, Henry Schreiber, Chengxin Zhang, Quancheng Liu, Scott Hultgren, Lydia Freddolino

## Abstract

The rapid advancement of high-throughput sequencing technologies has vastly increased the number of known protein sequences, but the experimental characterization of their structures and functions lags behind. This gap in knowledge impedes our understanding of biological mechanisms of these proteins, hinders the interpretation of high-throughput experiments, and exposes a significant challenge in modern biology: deducing the structural and functional information of proteins based on their sequences. Most computational approaches rely on homology with well-annotated proteins, yet many proteins lack identifiable homologues, reducing the power of this approach. Here, we integrated cutting-edge protein structure and function prediction methods to develop a complete sequence-structure-function pipeline that predicts structures and functions based on primary sequences. We applied this pipeline to predict the structure and function of all proteins in *Escherichia coli* UTI89, a model strain of uropathogenic *E. coli*. Based on the predicted functions, we performed enrichment analysis on the whole genome and revealed the possible roles and related biological mechanisms of poorly annotated proteins in this organism. Moreover, the performance of our pipeline was further validated through detailed case studies of the *UTI89_C0931* and *ybtS* genes. Finally, we compiled the UTI89 structure and function database (https://seq2fun.dcmb.med.umich.edu/UTI89), offering it as a community resource to aid researchers in elucidating the roles of unannotated proteins in uropathogenic *E. coli*. This database aims to bridge critical knowledge gaps in microbial pathogenicity and resistance, enhancing our capacity to tackle emerging health threats.

## Introduction

Over the past few decades, advancements and rapid developments in sequencing technologies have enabled the processing of longer read lengths and substantially reduced the costs of sequencing, thus making genome sequencing affordable [1, 2]. Consequently, there has been an explosive increase in the availability of genomic and protein sequences. The total number of protein entries in the automated TrEMBL database has experienced an exponential increase, surpassing 253 million entries as of November 2024 [3]; however, the expansion of Swiss-Prot, a manually curated database dedicated to protein sequences with reliable annotations, has significantly stagnated, and less than 0.5% of known protein sequences currently are associated with curated Swiss-Prot annotations [3]. In addition, many of the proteins in the Swiss-Prot database remain uncharacterized. Even for the well-studied *Escherichia coli* K12 strain MG1655 [4], more than 30% of genes remain uncharacterized or poorly annotated (annotation score ≤2 according to the UniProt database). The proportion of uncharacterized proteins is even higher in less-studied organisms [5]. Nevertheless, these poorly annotated proteins may play an important role in physiological processes; transposon library and dCas9 screens, for example, routinely identify strong fitness effects from knockdown/deletion of poorly annotated proteins [6, 7]. Therefore, a comprehensive understanding of protein structures and functions, particularly those of poorly annotated proteins, is crucial for a thorough grasp of the biological mechanisms of the entire genome.

Uropathogenic *Escherichia coli* (UPEC) are the primary pathogens responsible for urinary tract infections (UTIs) [8], contributing to around 75% of uncomplicated cases and over 50% of complicated cases [9]. UTIs are among the most common bacterial infectious diseases afflicting humans [10], with approximately 150 million cases reported annually [11]. Consequently, they pose a significant issue for healthcare systems and result in billions of dollars of financial burden including direct and indirect costs [12]. UPEC can reside within the gastrointestinal tract asymptomatically, but can opportunistically colonize the urinary tract and, during ascending infections, cause cystitis (bladder infections) or pyelonephritis (kidney infections). In recent years, UPEC strains have exhibited a startling rise in resistance to many antimicrobial treatments [11]. As a result, there is an urgent need to identify new antimicrobial targets for the treatment of UTIs and to determine potential vaccine targets to prevent infection [12, 13]. To achieve this, we must comprehensively understand the structure and function of bacterial genes essential for colonization, growth, and survival of UPEC during UTIs. UTI89, classified under multilocus sequence type 95 (ST95), is a model strain of UPEC [11] that has been used to extensively characterize host and pathogen outcomes to UTI in mouse models [14–20] and was originally isolated from a patient with an acute bladder infection [5]. It is frequently utilized in studies investigating virulence genes associated with UTIs [5, 11, 21]. Yet, based on UniProt, more than 88% (4549 out of 5192) proteins in the UTI89 genome remain poorly annotated.

In this study, we developed an automated hybrid pipeline for genome-wide protein structure and function prediction, which we then applied to the entire UTI89 proteome (although the same workflow should be widely applicable to other organisms). The pipeline comprises two components: the first module, sequence-to-structure module, employs a variety of protein structure prediction approaches including LOMETS3 [22], MODELLER [23, 24], D-I-TASSER [25], DMFold [26], and models from the protein databank (PDB) [27] and AlphaFold database (AFDB) [28, 29] to obtain structures or structural models of all input protein sequences. The second module, structure-to-function, utilizes our recently developed protein function prediction tool, StarFunc [30], to predict the functions of these proteins. StarFunc comprehensively utilizes sequence homology information, structural information, protein interaction information, and information inferred by deep learning to predict protein function, which overcomes defects of protein function prediction methods that only rely on homology transfer. Based on our predictions, we provide i) more comprehensive and enriched functional annotations than UniProt and ii) more accurate structural predictions of uncharacterized proteins than AFDB through our hybrid pipeline. First, we provide 31,512 additional GO term annotations, resulting in increases of 15.0%, 64.1%, and 31.1% in the ‘molecular function’, ‘cellular component’, and ‘biological process’ aspects, respectively. For proteins lacking experimentally determined structures and with AFDB models with low confidence (i.e., having predicted local distance difference test (pLDDT) scores below 70), our pipeline offers more reliable structural predictions, with average pLDDT improved by at least 15.9% and newly available models for 13 proteins (average pLDDT=82.79). Based on our results, we examined features of the UTI89 proteome more deeply. Here, GO term enrichment analysis revealed that some key functional categories, notably including bacteria-host and phage-bacteria interactions, are systematically missed in the previously existing UniProt-derived dataset but significantly overrepresented among poorly annotated proteins, with several findings supported by existing literature. Further, case studies on *UTI89_C0931* and *ybtS* genes further validated the reliability of our predictions. To support broader research efforts, we compiled all structure and function predictions into a centralized, publicly available database. This sequence-structure-function pipeline, together with future improvements, is publicly available and will be extended to other less-studied organisms to advance the functional characterization of poorly annotated proteins and provide valuable resources for the biomedical research community.

## Materials and methods

In this section, we introduce our automated sequence-structure-function pipeline (illustrated in **Figure 1**), which is composed of two main components. The first component involves generating a three-dimensional protein structure from the input amino acid sequence, representing the sequence-to-structure process. The second component utilizes StarFunc [30] to predict the corresponding protein function based on the derived structure.

**Figure 1.**
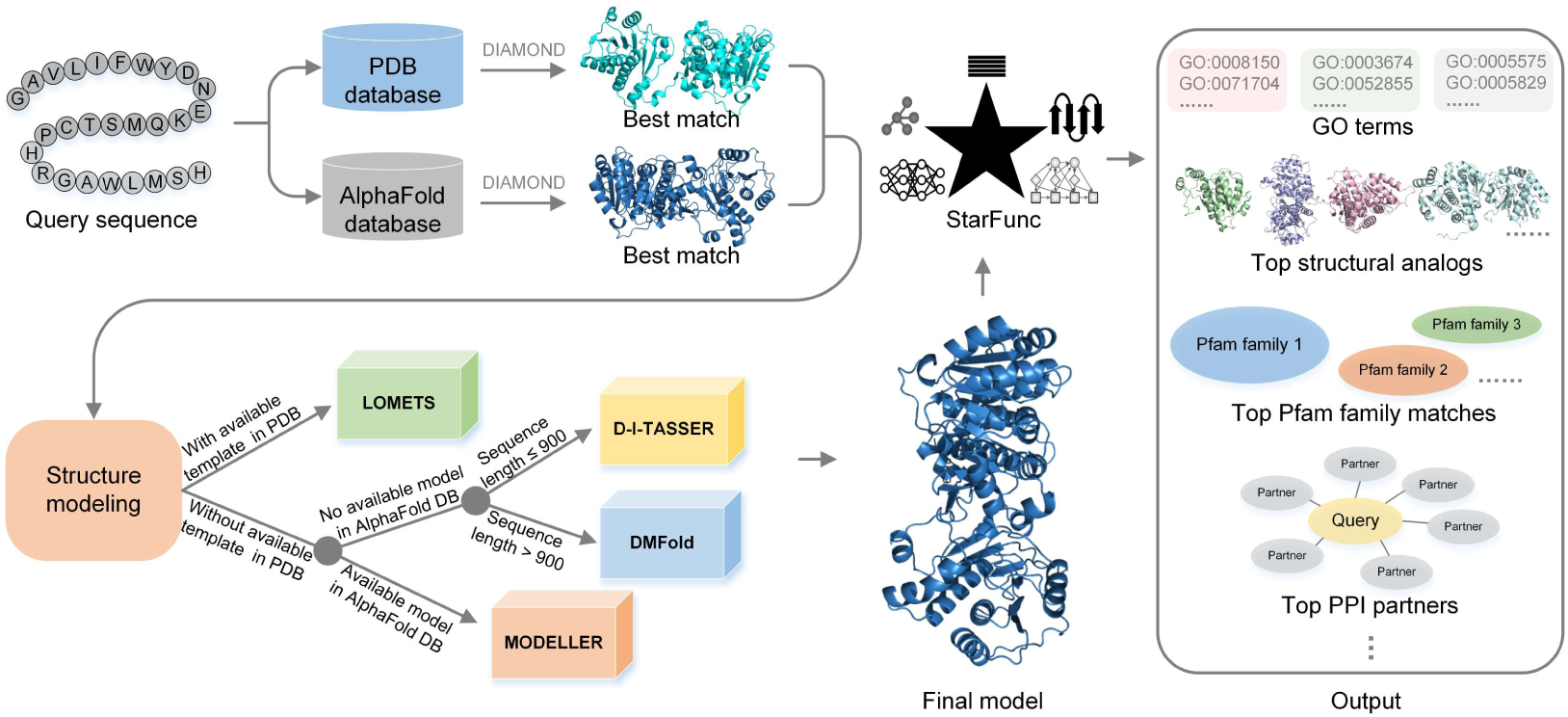
Sequence-structure-function pipeline for whole-proteome structure and function prediction. This pipeline features two modules: a protein structure generation module and a protein function prediction module. The structure generation module integrates LOMETS3 [22], MODELLER [23, 24], D-I-TASSER [31], and DMFold [26], modeling structures based on the quality of templates and AlphaFold2 models identified by DIAMOND [32, 33]. If an identical structure already exists in the PDB, the experimentally determined structure is used directly. Subsequently, the function prediction module utilizes StarFunc to predict protein functions [30].

### Protein structure prediction

We obtained structural models of all 5,192 proteins in the *E. coli* UTI89 genome (UniProt reference proteome ID UP000001952 [5]) using a hybrid protein structure prediction pipeline, primarily consisting of four methods: MODELLER [23, 24], LOMETS3 [22], D-I-TASSER [25], and DMFold [26], as shown in **Figure 1**. The workflow of the protein structure prediction pipeline is as follows: (1) For each input protein sequence, DIAMOND [32, 33] with default parameters is used to identify homologous templates from the PDB [34] and AFDB [28], and the template with minimum e-value is selected from each database. (2) If a structure from the PDB exactly matches the query sequence and contains no missing residues, it is directly adopted as the final model. (3) If no perfect match is available, but the sequence identity between the PDB-derived template and the query is ≥0.95, the final structure is generated using LOMETS3 based on the template. (4) If the above conditions were not met, a template from the AFDB was considered. If the AFDB-derived template had an identical sequence to the input and an average predicted pLDDT ≥70, it was directly used. (5) If the AFDB template has a sequence identity in the range [0.95, 1.0) and an average pLDDT ≥70, MODELLER is used to refine the template to generate the final structure. (6) In all other cases, the final model is predicted using either D-I-TASSER (for sequences <900 amino acids) or DMFold (for longer sequences).

### Structure-based protein function prediction

Protein functions were predicted using StarFunc [30], our most recent protein function prediction approach. StarFunc was run with default settings, and function predictions with a confidence score (Cscore) >0.5 were considered confident, consistent with our previous work on JCVI-syn3.0 [35]. A higher Cscore indicates greater confidence in the predicted function.

### Enrichment Analysis

To investigate which functions are commonly associated with poorly annotated (annotation score ≤2 in UniProt) versus well annotated (annotation score ≥3 in UniProt) proteins, a Chi-squared test with Benjamini-Hochberg FDR correction [36] was first performed to assess whether the occurrence frequency of each GO term across the entire *E. coli* UTI89 genome differs significantly between these two groups (statistical calculations were performed using appropriate scipy [37] functions unless otherwise noted). The higher annotation scores indicate greater reliability and specificity of the function annotation for a given protein. For each GO term exhibiting a significant difference, its enrichment in poorly annotated proteins was quantified using the log odds-ratio of the representation of that GO term *q* in poorly annotated proteins relative to well-annotated proteins, defined as follows:

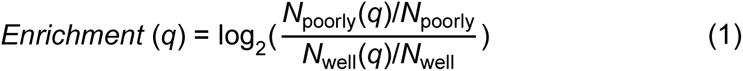

where *N*_poorly_ and *N*_well_ represent the number of poorly and well-annotated proteins, respectively; *N*_poorly_(*q*) and *N*_well_(*q*) are the number of poorly and well-annotated proteins with GO term *q*, respectively. Since the argument of the logarithm is in the form of a ratio between two rates, a p-value from the rate ratio test can be calculated for each enrichment score [38]. Due to the directed acyclic graph architecture of GO terms, if a child term *q* is associated with a protein, all its parent terms will also, by definition, be associated with this protein, and in some cases the set of proteins in our data set with a particular child term and a corresponding parent term will be identical. To reduce such redundancy in enrichment analysis, parent terms were excluded if they had the same enrichment and *p*-values as their corresponding child terms; *p*-values were subsequently FDR corrected using the Benjamini-Hochberg method.

Notably, some GO terms, such as “GO:0008152/metabolic process”, are too general to indicate a specific function. Consequently, similar to our previous whole proteome function-prediction study on JCVI-syn3.0 [35], these common GO terms were excluded from all steps of our analysis based on the following criteria. For biological process (BP) and molecular function (MF) terms, any predicted GO terms associated with more than 25% of UTI89 proteins were removed. For cellular component (CC) terms, any GO terms associated with more than 65% of proteins were removed. This is because *E. coli*, as a prokaryote, has a less complex cellular architecture compared to eukaryotes, limiting the specificity of GO terms that can describe its CC. As a result, a total of 17 common GO terms were excluded: 6 for BP, 8 for MF, and 3 for CC, respectively. Details of these 17 GO terms can be found in **Supplementary Table S1**. Besides the removed common GO terms, the remaining GO terms can be considered as specific GO terms. In addition, some GO terms that are obviously incorrect for *E. coli*, such as “GO:0005739/mitochondrion” and its children terms, were removed from the GO terms predicted by StarFunc, reflecting a common phenomenon of taxonomically obvious mis-annotations [38]. A total of 4 GO terms were excluded for taxonomic reasons, with details provided in **Supplementary Table S2**.

### Benchmarking datasets

To evaluate the performance of our workflow in protein function prediction and to investigate the relationship between the predicted structure quality (pLDDT) and StarFunc’s prediction performance, we constructed two independent non-redundant test datasets, referred to as **Dataset I** and **Dataset II**. Detailed procedures for constructing **Dataset I** and **Dataset II** are provided in **Supplementary Text S1**. **Dataset I** was curated to avoid data leakage and redundancy, whereas **Dataset II** was constructed to ensure that each protein is associated with models spanning distinct pLDDT ranges.

**Dataset I**, intended to test the overall performance of function prediction, comprises 250 proteins with new annotations since all computational pipelines were developed. This dataset was used to evaluate and compare the performance of our pipeline with two widely used methods: a homology-based function transfer approach (represented by ‘sequence-based’) and a representative pure deep learning method, DeepGOPlus [39]. The details of sequence-based method used in this study can be found in **Supplementary Text S2**. The performances of these methods were evaluated using the maximum of the information accretion-weighted F-measure (wF_max_, details can be found in **Supplementary Text S3**), as adopted in CAFA [40]. The higher of wF_max_ represents the better performance.

**Dataset II**, intended to assess the effects of model quality on the accuracy of our functional predictions, consists of 128 proteins; for each such protein, we obtained 1,000 structure models by running AlphaFold2 200 times with different random seeds (100 runs with default settings and 100 runs without using templates and MSAs). The pLDDT is a per-residue measure of local confidence; it is scaled from 0 to 100, with higher scores indicating higher confidence and usually a more accurate prediction. We then stratified the predicted structures across three distinct pLDDT intervals: (0, 50], (50, 70], and (70, 100]; for each protein, we separately selected the model with the median pLDDT from each interval as input for StarFunc.

### UTI89 biofilm formation experiments

#### Bacterial strains

Deletion mutations of a clinical UPEC isolate, UTI89 [7], were made using the lambda red recombinase method, as previously described [8], using pKD4 or pKD13 as a template and the primers as listed in **Supplementary Table S3**. PCR was performed with flanking primers to confirm the appropriate deletions. Antibiotic insertions were removed by transforming the mutant strains with pCP20 [9] expressing the FLP recombinase. The resultant strains subsequently had no additional antibiotic resistance compared with the parental WT strain. A complete list of bacterial strains used in this study is given in **Supplementary Table S4**.

#### Bacterial cultures

For biofilm assays, bacteria were grown in LB or YESCA media in wells of PVC microtiter plates. For LB and YESCA biofilms, after 48 hours of growth, wells were rinsed and stained with crystal violet for quantification as described [10]. Pellicle was assessed by inspection at 48 or 72 hours for the presence or absence of a coherent surface membrane at or near the air-liquid interface. A pellicle exhibits sufficient mechanical integrity that it can be lifted intact from the culture using a forceps.

#### Mass spectrometry

Metabolites were extracted by applying 1 mL of conditioned media to a C18 solid phase extraction cartridge (100 mg, Waters) conditioned with acetonitrile and water, respectively. After washing with 1 mL water, retained metabolites were eluted with 1 mL acetonitrile and the resulting eluate was concentrated under a stream of dry nitrogen. Untargeted profiling of the extracellular metabolome of pellicle cultures was performed using an AB Sciex QTrap 4000 mass spectrometer (Foster City, CA) coupled to a Shimadzu LC20-AD XR HPLC equipped with a Supelco C18 column (1×150mm). The flow rate was 200 µL/min with a gradient as follows: Solvent A (0.1% formic acid) was held constant at 98% for 2 minutes, increased to 58% solvent B (0.1% formic acid in acetonitrile) in the next 10 minutes, and then to a 98% B in the next 2 minutes. The column was re-equilibrated for 3 minutes. Ionization was in positive ESI mode. The spray voltage on the mass spectrometer was held constant at 5.0 kV and the temperature was 500. The mass spectrometer was operated in the linear ion trap mode with scanning from m/z 50 to 2000. Comparison of major components was achieved by manual inspection of total ion chromatograms for differentially expressed chromatographic peaks and inspection of the representative mass spectra.

#### Preparation of unlabeled and deuterated ferric yersiniabactin

Unlabeled yersiniabactin was produced by inoculating M63 minimal media containing 10 mg/L niacin and 0.2% glycerol with 20 mL of UTI89Δ*entB*, growing in shaking conditions at 37 C for 18 hours. Cells were removed by centrifugation. 3.75 mmol/L of ferric chloride was added to the culture supernatant to convert yersiniabactin to its ferric complex. The resulting precipitate was removed by filtration through a 0.22 µm filter. Ferric yersiniabactin was concentrated by adsorption to C18 silica followed by elution with 80% methanol. A red-orange peak with a visible spectrum absorbance maximum at 385 nm was collected, dried in a centrifugal evaporator, and confirmed as ferric yersiniabactin using liquid chromatography-mass spectrometry as described [11]. Concentration was determined by absorbance at 385 nm [12].

#### Yersiniabactin quantification

Absolute quantification of yersiniabactin content in culture media was determined by stable isotope dilution mass spectrometry using d4-ferric yersiniabactin as the internal standard and comparing to a standard curve. To each culture supernatant, d4-ferric yersiniabactin standard was added followed by 3.75 mmol/L of ferric chloride. Supernatants were then fractionated by C18 solid phase extraction (100 mg, Waters) by washing with 20% methanol and eluting with 80% methanol. Eluates were concentrated in vacuo and the ratio of unlabeled to labeled yersiniabactin was determined as previously described [11]. The precursor ion for MS/MS analysis of d4-ferric yersiniabactin is m/z 539 and the product ions are m/z 353 and 188.

## Results and Discussion

### General statistical analysis of function annotations

We applied our automated sequence-structure-function pipeline to predict the structures and functions of the entire *E. coli* UTI89 proteome and compared these predictions with the function annotations available in UniProt (**Figure 2**).

**Figure 2.**
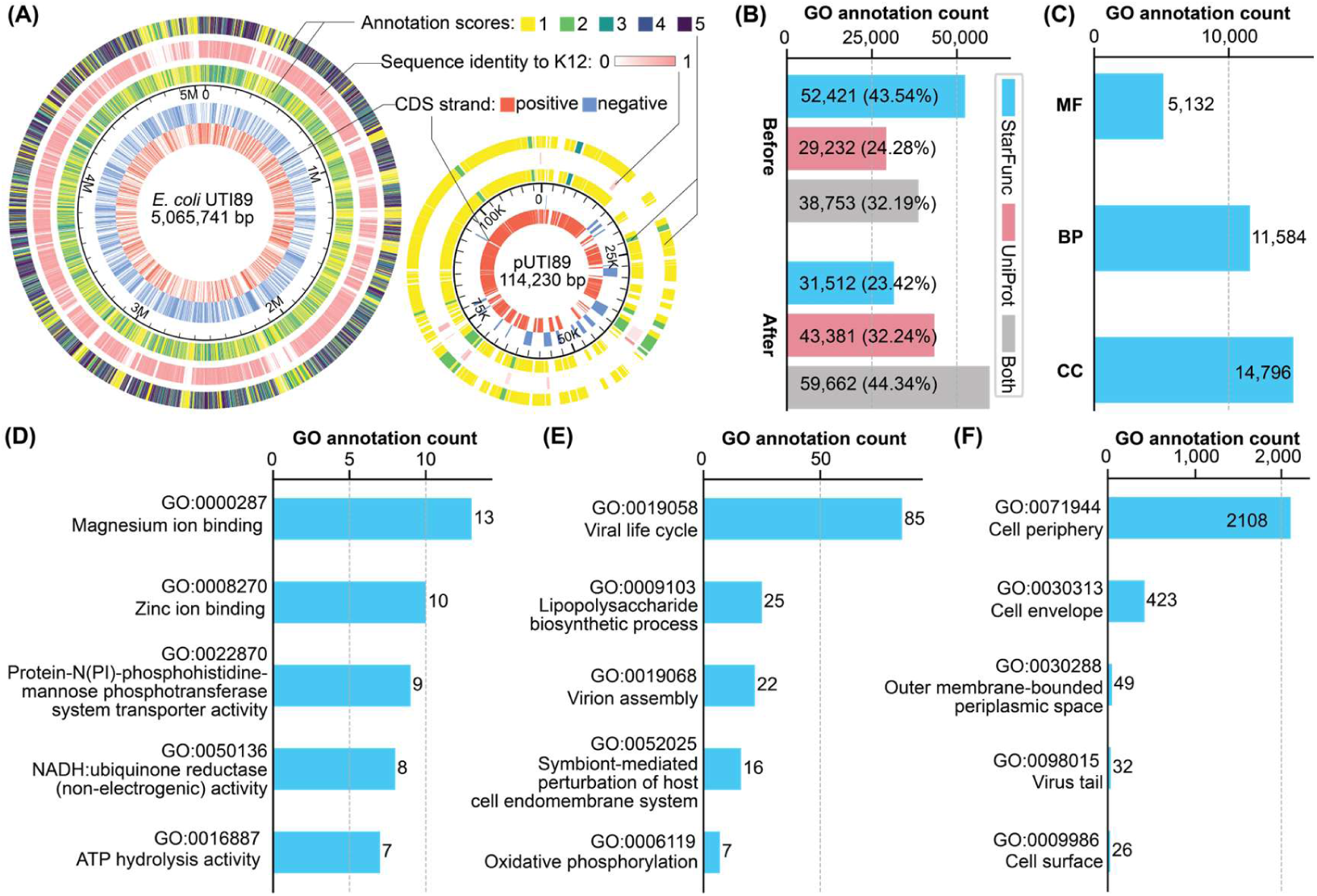
Statistics for protein function annotations in the *E. coli* UTI89 proteome. **(A)** The distributions of coding strand orientation (1st and 2nd rings), annotation scores (3rd and 5th rings; before and after the *E. coli* K12-based update, respectively), and maximum sequence identity to K12 proteins (4th ring) are shown for each protein in the UTI89 proteome. **(B)** The number of GO term annotations predicted by StarFunc, provided only by UniProt, or shared by both sources. “Before” refers to the GO term annotations originally provided by UniProt for *E. coli* UTI89, whereas “After” refers to the updated annotations obtained by incorporating transferable GO terms from *E. coli* K12, also sourced from UniProt. **(C)** The number of new GO term annotations only provided by StarFunc for three different aspects of protein function. **(D)**-**(F)** Top 5 GO terms with the highest number among all new GO terms, **(D)** for molecular function (MF), **(E)** for biological process (BP) and **(F)** for cellular component (CC).

According to the raw data provided by UniProt, out of the 5,192 proteins in the UTI89 proteome, only 4 proteins were assigned an annotation score of 5 (<0.1%), 39 proteins were assigned a score of 4 (0.8%), 600 proteins were assigned a score of 3 (11.6%), 1,408 proteins had a score of 2 (27.1%), and 3,141 proteins were assigned a score of 1 (60.5%), as shown in the 3rd ring in **Figure 2A**. Here, higher annotation scores indicate greater reliability and specificity of the function annotation for a given protein. An annotation score of 5 represents the best-annotated entry, while a score of 1 signifies entries with only basic annotation details. Therefore, unless otherwise stated, we use the phrase ‘poorly annotated’ to refer to proteins with UniProt annotation scores of 1 or 2, and ‘well annotated’ for proteins with annotation scores 3 or higher in this study. Poorly annotated proteins typically lack functional homologs or have homologs without consistent functional annotations, whereas well-annotated proteins often match homologs with related functions or have been extensively studied. *E. coli* K12 is one of the most extensively studied and commonly used laboratory strains, sharing many identical or highly similar genes with UTI89. Leveraging the well-characterized *E. coli* K12 proteome, we transferred annotations of homologous proteins to the UTI89 proteome to construct a more robust and comprehensive functional annotation set. Therefore, MMseqs2 [41] and CD-HIT [42] were used to identify proteins in the K12 proteome with high sequence identity (>90%) to those in the UTI89 proteome. For the same target sequence, the higher sequence identity between MMseqs2 and CD-HIT is taken as the final sequence identity between the query sequence and the target sequence. For UTI89 proteins with low annotation scores, the annotation information, including annotation scores and GO terms, was updated by K12 proteins with the highest annotation score when available. The updated annotation scores for the UTI89 proteome are summarized in the 5th ring in **Figure 2A**. The number of proteins with an annotation score of 5 increased from 4 (<0.1%) to 1,472 (28.4%), and those with a score of 4 increased from 39 (0.8%) to 613 (11.8%). The number of proteins with a score of 2 decreased from 1,408 (27.1%) to 662 (12.8%), and proteins with a score of 1 dropped from 3,141 (60.5%) to 1,879 (36.2%). Nonetheless, 48.9% of the proteins (2,541) remain poorly annotated. Moreover, as clearly shown in **Figure 2A**, the majority of proteins (∼99.3%) encoded on the large virulence plasmid in *E. coli* UTI89 are poorly annotated. Uncovering the potential functions of these poorly annotated proteins is crucial for understanding the pathogenicity of *E. coli* UTI89 and for improving the treatment of urinary tract infections.

Although annotation transfer from *E. coli* K12 substantially improved the overall annotation scores of the UTI89 proteome, many proteins still lacked detailed functional descriptions, particularly in terms of GO term assignments. To address this limitation, we used the pipeline to systematically predict and expand the structures and GO annotations for the UTI89 proteome. On the *E. coli* UTI89 proteome, a total of 91,174 specific GO term annotations was predicted using the pipeline. In comparison with the original function annotations provided by UniProt, 29,232 specific GO annotations were unique to UniProt, 38,753 were shared between UniProt and StarFunc, and 52,421 specific GO annotations were provided only by StarFunc (**Figure 2B**). Here, the GO terms provided by UniProt are from the UniProt-GOA project [43]. Even after updating the function annotations of UTI89 proteome using GO terms from highly similar and better-annotated proteins in the K12 strain, StarFunc still can provide 31,512 additional GO term annotations (**Figure 2B**). Among these GO annotations, 5,132 pertained to MF, 11,584 to BP, and 14,796 to CC (**Figure 2C**). For each aspect, the top 5 new GO terms with the highest number are shown in **Figures 1D**-**F**. Using the pipeline, we identified an additional 25, 22, 16, and 9 UTI89 proteins potentially involved in “lipopolysaccharide biosynthesis/GO:0009103”, “symbiont-mediated perturbation of the host endomembrane system/GO:0052025”, and “mannose PTS transporter activity /GO:0022870”, respectively. The predictions not only supplement existing function annotations but also reveal uncharacterized proteins that may play crucial roles in UTI89’s metabolism, environmental adaptation, and virulence.

The UTI89 genome, like many bacterial genomes [44, 45], contains a fraction of proteins annotated with functions related to the “viral life cycle/GO:0019058” and “virion assembly/GO:0019068”, likely reflecting more detailed annotations of genes from integrated prophages. Using our pipeline, we significantly expanded the annotation of genes related to the viral life cycle and virion assembly in the UTI89 genome. StarFunc uniquely annotated 85 proteins with the GO term “viral life cycle”, among which 22 were further annotated with “virion assembly”. Among these 85 proteins, 65 (76.5%) were identified as phage-related genes by PHASTER [46], based on a criterion of greater than 90% overlap with predicted prophage (see **Supplementary Table S5** for details). In particular, functions associated with cell periphery and virus tail have not been recorded for many of the corresponding proteins in UniProt annotations, but additional proteins were identified with these two functions by using the pipeline. The more detailed annotation of prophage-associated genes provides a powerful example of the utility of our global annotation pipeline; while a detailed manual investigation of many of these cases would likely lead a researcher to similar conclusions regarding the functions of the prophage-associated genes identified here, the advantage of our approach is that it provides automated supplementation of available UniProt annotations that are of particular utility alongside high throughput studies such as genetic screens.

### Structure-based function prediction increases the coverage of function annotation

According to the most recent the 5th Critical Assessment of Function Annotation (CAFA5) challenge, StarFunc demonstrated its advantage, ranking 5th out of 1,625 teams from 96 countries [30]. Here, we further evaluated the performance of our pipeline using an independent test set of 250 newly released proteins (see **Dataset I** for details). We benchmarked it against a widely used homology-based function transfer method (represented by ‘sequence-based’; details in **Supplementary Text S2**) and a representative deep learning approach (DeepGOPlus) [39] to highlight the overall performance of the full pipeline. Their predictive accuracy was assessed using wF_max_, which measures the overall balance between precision and recall across all confidence thresholds, providing a robust estimate of global prediction accuracy for functional annotations. We next investigated how many UTI89 proteins could be assigned specific GO term annotations. Here, UTI89 proteins were categorized based on their UniProt annotation scores (including K12-based updates and original annotations).

Overall, for the three GO aspects, our pipeline demonstrates superior performance in BP and CC compared to the better of the two baseline methods (**Figure 3A**), achieving a 2.3% higher wF_max_ for BP (0.445 vs. 0.435 for the sequence-based method; p=2.46×10^−39^, Wilcoxon signed rank test, which is also used throughout this work for similar comparisons unless otherwise noted) and an 8.7% improvement for CC (0.663 vs. 0.610 for DeepGOPlus; p=0.00). For MF, although the difference is statistically significant (p=1.24×10^−5^), both our pipeline and the sequence-based method achieve an identical average wF_max_ of 0.499, indicating comparable predictive performance in practice. Nevertheless, when averaging across the three GO aspects, our pipeline attains an average wF_max_ of 0.535, which is 4.7% and 5.9% higher than that of the ‘sequence-based’ (0.511) and DeepGOPlus (0.505), respectively. These results further highlight the overall robustness and effectiveness of our pipeline, which delivers consistently strong performance across GO aspects and achieves higher average functional annotation accuracy than both homology-based and deep learning-based methods.

**Figure 3.**
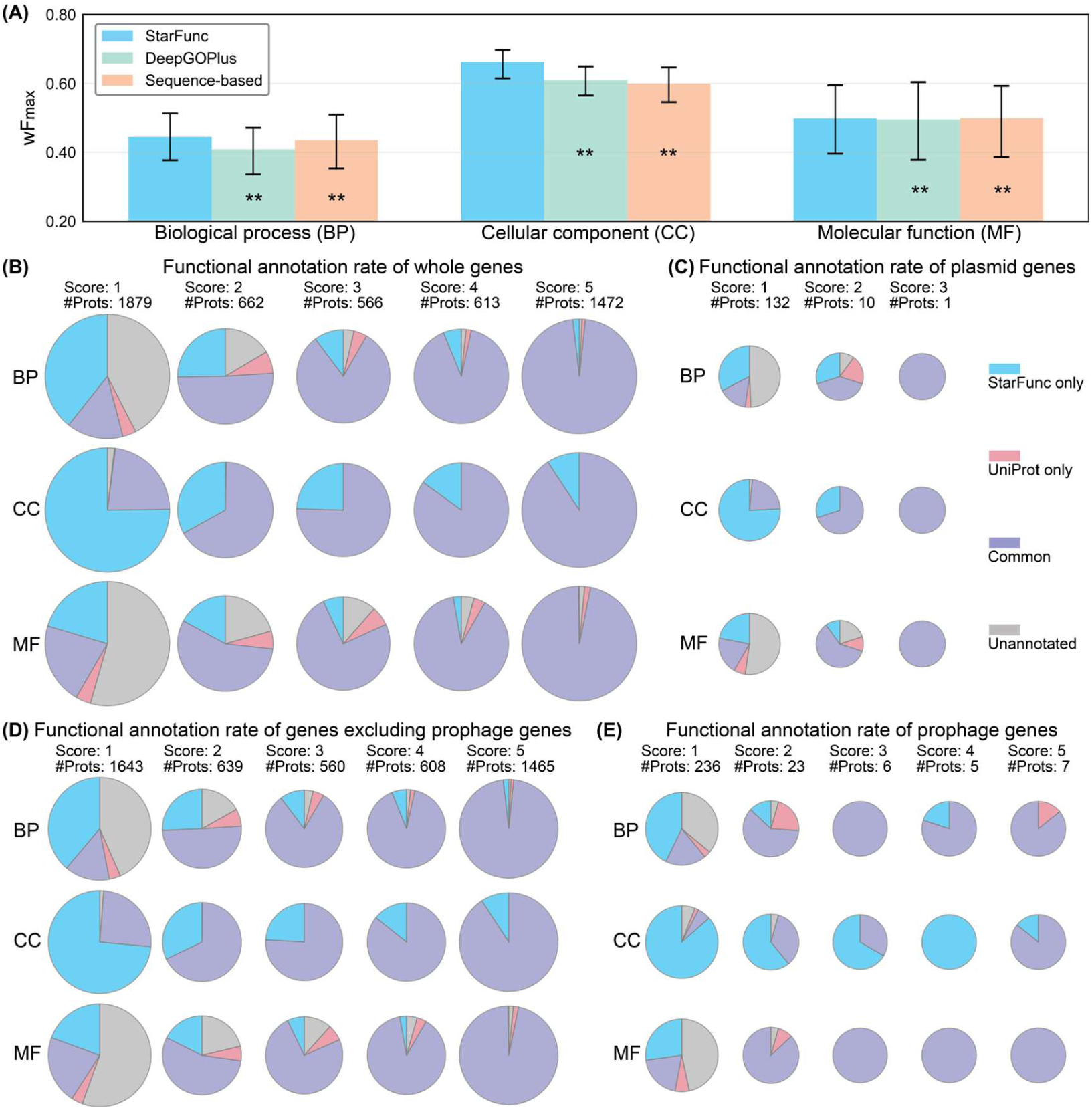
Overall performance of sequence-structure-function pipeline with widely used methods, improving the coverage of protein function annotations in UniProt. **(A)** for the three different aspects of GO, the wF_max_ of StarFunc, DeepGOPlus, and a homology-based function transfer method (represented by ‘sequence-based’) on our set of 250 test proteins. 95% confidence intervals were estimated using 10,000-resample bootstrap via scipy.stats.bootstrap, accounting for limited sample size. Asterisks denote statistical significance levels: * indicates p <0.05 and ** indicates p <0.01. **(B)**-**(E)** Functional annotation rates of UTI89 proteins across annotation score categories, **(B)** for whole UTI89 genome, **(C)** for plasmid-encoded genes, **(D)** for genes excluding phage-derived sequences, and **(E)** for phage-derived genes. Pie charts show the proportion of proteins with at least one specific GO term assigned in the three GO aspects. Proteins were classified into five categories (Score 1-5) based on their UniProt annotation score, with the total number of proteins in each category indicated above (#Prots). Blue and purple segments represent proteins annotated by the proposed pipeline, red and purple represent proteins annotated by UniProt, and grey indicates unannotated proteins. Overlap between the two methods is shown in purple. The specific numbers can be found in **Supplementary Tables S6-S17**.

Based on the predictions from our pipeline and the annotations from UniProt, we counted how many UTI89 proteins (now considering the entire proteome) could be assigned at least one specific GO term by each method. The broader coverage of our pipeline is particularly evident for the poorly annotated proteins (**Figures 3B**). Even for these proteins with an annotation score of 3, our pipeline still can assign specific GO terms to more proteins than UniProt. The broader coverage of our pipeline is most evident for proteins with an annotation score of 1. For instance, the pipeline annotates 54.0%, 97.9%, and 41.7% of these proteins with at least one specific BP, CC, and MF terms, respectively, which are 35.8%, 75.0%, and 16.4% higher than that of UniProt. Especially for CC terms, almost 100% of proteins were assigned specific GO terms by our pipeline. Unsurprisingly, for proteins with annotation scores of 4 or 5 in UniProt, the differences in coverage between existing UniProt annotations and those derived from our pipeline are minimal. Notably, combining UniProt GO terms with those predicted by StarFunc led to consistent improvements in the percentage of annotated proteins across all protein sets. The greatest increase, reaching 75.0%, was observed for the CC aspect of proteins with an annotation score of 1. This is because StarFunc integrates not only annotations from sequence- and structure-similar proteins, but also incorporates protein-protein interaction-based, Pfam-based, and deep learning-based annotations, thereby providing a more comprehensive set of GO annotations compared to UniProt. Even for proteins with the highest annotation scores, StarFunc is capable of providing additional, functionally specific GO terms. For example, the hydrogenase *HybC* (UniProt accession Q1R701), a core component of the *E. coli* [NiFe]-hydrogenase isoenzyme 2 (Hyd-2), lacked specific BP GO terms despite being annotated by transfer from a well-studied homolog in *E. coli* K12 (P0ACE0, annotation score 5). Using our prediction pipeline, we assigned two highly relevant BP GO terms to *HybC*: “anaerobic respiration/GO:0009061” and “respiratory electron transport chain/GO:0022904”. These predicted functions align well with previous biochemical studies that have characterized Hyd-2 as an O_2_-sensitive [NiFe]-hydrogenase complex involved in hydrogen uptake and electron transfer under anaerobic respiratory conditions [47]. The prediction that UTI89 encodes anaerobic respiration-related proteins such as *HybC* provides insight into how the pathogen may adapt to oxygen-limited environments in the urinary tract, thereby enhancing its colonization and infection efficiency. This also suggests potential avenues for experimental validation and therapeutic intervention. In addition, this also shows the pipeline’s utility in discovering potential functions and providing valuable candidates for further experimental validation (and, importantly, providing those annotations without manual intervention).

We further assessed the annotation coverage of protein-coding genes across three distinct subsets of the *E. coli* UTI89 genome on different annotation scores. These subsets include: 143 genes located on plasmid (**Figure 3C**); 4,915 chromosomal genes excluding phage-derived sequences (**Figure 3D**); 277 phage-derived genes (**Figure 3E**). The results indicate that StarFunc demonstrates a pronounced advantage in annotating poorly characterized proteins. For plasmid-encoded proteins, 99% are poorly annotated, and none received an annotation score above 4, indicating that plasmid genes remain understudied and limited annotations were provided in UniProt. By contrast, StarFunc improves the functional annotation of these poorly annotated proteins, assigning at least one specific GO term in the BP, CC, and MF aspects to 47.8%, 98.5%, and 41.7% of proteins with an annotation score of 1, respectively. Across plasmid, chromosomal, and phage-derived genes, StarFunc substantially expands GO term coverage beyond what is provided by UniProt. These findings further highlight the utility of StarFunc in extending and complementing existing annotations, particularly for genes of pathogenic relevance, where its predictions may yield novel biological insights.

Overall, our pipeline can provide more specific GO terms than UniProt for the majority of proteins, particularly for those that are poorly annotated. Meanwhile, these results suggest that GO terms predicted by StarFunc complement those from UniProt and integrating both annotations can offer researchers a more comprehensive set of annotations for any target proteome of interest.

### A tiered pipeline provides higher-confidence models for difficult protein structural targets

The functions of proteins depend on their unique three-dimensional structures [48]. Understanding their structure can facilitate a mechanistic understanding of their functions [29], an assumption underlying the structure-based annotation module of StarFunc [30]. Likewise, the structural quality of the input models may affect the reliability of StarFunc’s GO term predictions. In this section, we first examined the relationship between model confidence, measured by pLDDT, and StarFunc performance, assessed by wF_max_. The higher pLDDT scores generally indicate greater structural accuracy. We then compared the pLDDT scores of models generated by our multi-tiered pipeline with those from uniform use of AlphaFold2 to evaluate whether our pipeline produces higher-confidence structures (**Figure 4**).

**Figure 4.**
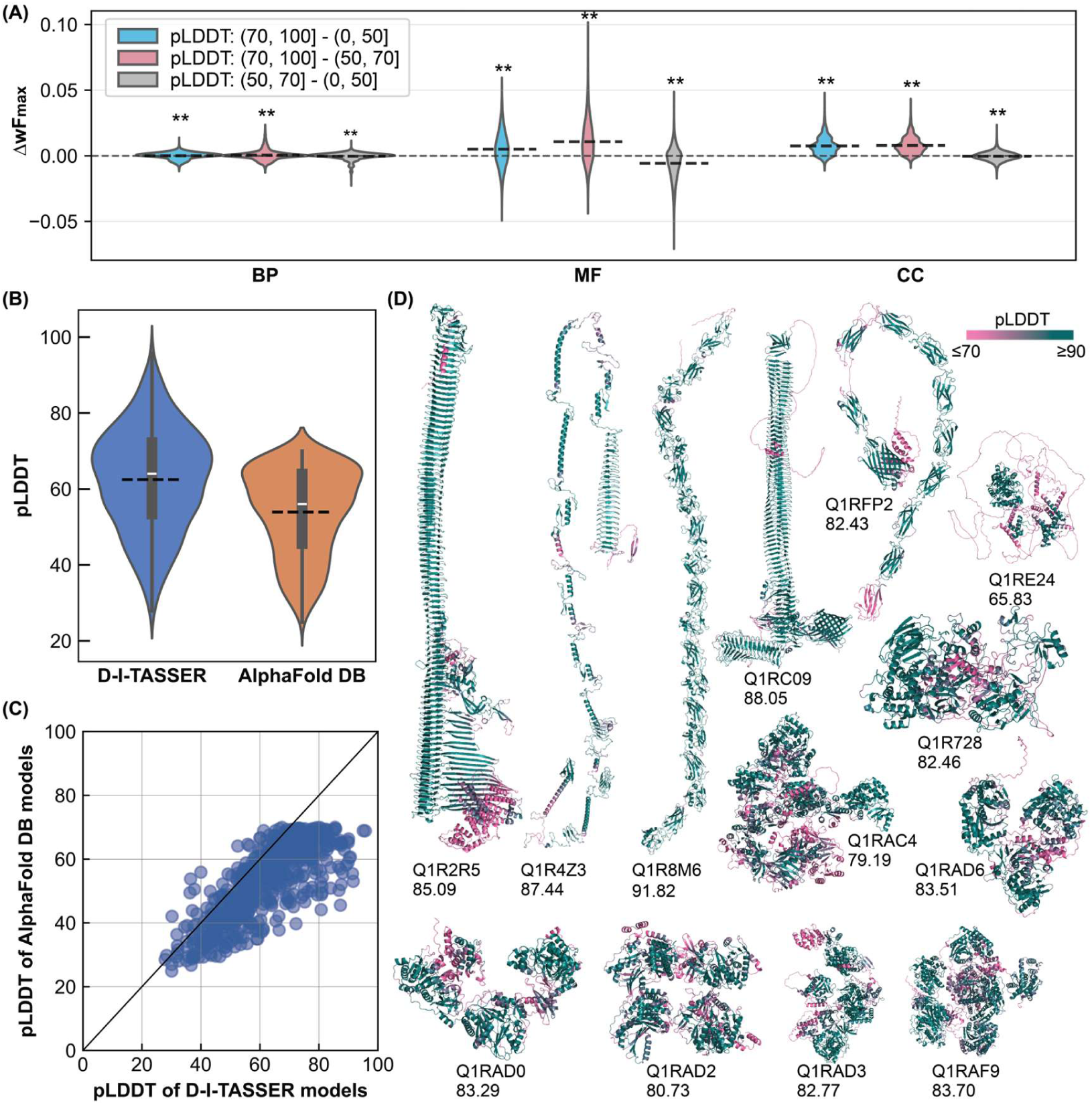
Impact of model confidence on StarFunc performance and comparison between D-I-TASSER and AFDB models. **(A)** Effect of models with different pLDDT scores on StarFunc GO term prediction performance. Asterisks denote statistical significance levels: * indicates p <0.05 and ** indicates p <0.01, and dash line denotes the mean ΔwF_max_. **(B)** Violin plot illustrating the distribution of pLDDTs for models generated by D-I-TASSER and from AlphaFold DB. The black dash line represents the average pLDDT and the white dot represents represent the median pLDDT among the 696 proteins. **(C)** Head-to-head comparison of pLDDTs between D-I-TASSER [31] and AlphaFold DB models. **(D)** DMFold [26] structure predictions on 13 proteins without any available structural information in UniProt, colored by pLDDT (pink, low; teal, high).

We evaluated the performance of StarFunc on **Dataset II**, which consists of 128 proteins, using AlphaFold2 models with varying pLDDT scores as input. We grouped the models into three pLDDT ranges (<50, 70-100, and 50-70) and compared their wF_max_. Since pLDDT scores above 70 generally indicate high-confidence structures [26], models with lower pLDDT were further divided into 50-70 and 0-50 to examine more precisely how model confidence affects the functional prediction performance of StarFunc. The results, shown in **Figure 4A**, indicate that across the three GO aspects, StarFunc achieves improved performance in MF and CC GO terms predictions when higher confidence models are used. For MF, models with pLDDT scores of 70-100 achieve a 0.9% higher wF_max_ than those with scores of 0-50 (0.542 vs. 0.537; p=9.01×10^−259^), and for CC, the increase is 1.2% (0.691 vs. 0.683; p=0.0). For BP, StarFunc achieves a comparable wF_max_ (∼0.432) regardless of the input model’s pLDDT. In retrospect, it is not surprising that higher pLDDT leads to more accurate prediction for MF and CC but not BP. Molecular functions such as catalytic activity and binding are often contingent upon specific shapes; specific structure features such as transmembrane helixes are indicative of cellular components; on the other hand, the relation between BP and structure is less obvious since many protein folds can often contribute to the same biological process. Thus we find that high-quality structural models, to some extent, enhance the accuracy of MF and CC predictions generated by StarFunc. Results from the StarFunc structure component also show a similar trend, in which higher confidence in the input structural models corresponds to better accuracy in functional prediction, as shown in **Supplementary Figure S1**.

Within the UTI89 proteome, 714 proteins lack high-quality homologous templates in the PDB and do not have high-confidence models available in the AFDB. Thus, for these proteins, we used D-I-TASSER (for 696 proteins) and DMFold (for 18 proteins) to predict their structures. D-I-TASSER and DMFold participated in the 15th and 16th Critical Assessment of Structure Prediction (CASP15 and CASP16), achieving state-of-the-art performance in protein structure prediction. D-I-TASSER ranked first in the monomer prediction category at CASP15 [31], while DMFold achieved top performance in complex structure prediction [26]. Notably, both methods (including the monomer version of DMFold) outperformed the publicly available AlphaFold2 and AlphaFold3 implementations in their respective categories [26]. Here, we compared pLDDT scores for 714 UTI89 protein models predicted by D-I-TASSER or DMFold with those from the AFDB to assess model reliability; among these, 696 proteins shorter than 900 amino acids were modeled using D-I-TASSER. The remaining 18 proteins, exceeding 900 amino acids, were predicted using DMFold, due to high resource consumption by D-I-TASSER for long proteins. Within this subset of 18 proteins, 13 lack AlphaFold2 models in UniProt. For 696 proteins with low-confidence models available in the AFDB, we compared the pLDDT scores of D-I-TASSER and AlphaFold2 predictions to assess whether our pipeline produces models with improved confidence. It is important to note that the model quality estimation systems of D-I-TASSER and AlphaFold2 are different. Therefore, we used the confidence scores of the top-ranked models from the intermediate stage of D-I-TASSER [31], which are also derived from the AlphaFold2 system, for fair and direct comparison with AlphaFold2 models. These intermediate models serve as a suitable basis for comparison because the final D-I-TASSER models are further optimized versions of these initial models, meaning that the final model predictions are likely to be more accurate than the intermediate models [31]. The mean and median pLDDTs of D-I-TASSER models are 62.47 and 64.00, respectively, representing a 15.9% and 14.3% improvement over the corresponding values for AlphaFold DB models (mean 53.92 and median 56.01, **Figure 4B**). D-I-TASSER generated models with statistically significantly higher pLDDT than AlphaFold DB (p=3.99×10^−95^), with 89.4% (622 out of 696) of the proteins having higher pLDDT in D-I-TASSER (**Figure 4C**).

For the other 18 proteins, DMFold was used to predict their structures due to their sizes exceeding 900 residues. Of these, only 5 proteins have structure models available in the AFDB, while the remaining 13 lack any structural information. The pLDDT details of these 18 proteins can be found in **Supplementary Tables S18 and S19**. For the 5 proteins, the average and median pLDDT scores of the DMFold models are 69.08 and 70.46, respectively, representing increases of 12.3% and 13.1% compared to those of the AFDB models. For other 13 proteins, the average and median pLDDTs of the DMFold models are 82.79 and 83.29, respectively, and their predicted structures are shown in **Figure 4C**. Notably, apart from the DNA translocase FtsK with UniProt accession Q1RE24, all remaining proteins have an average pLDDT exceeding 70. Thus, our pipeline is able to provide high-confidence models for the vast majority of UTI89 proteins, exceeding the accuracy and depth of existing databases and thus providing a richer collection of structural predictions than was previously available, and simultaneously providing a foundation for our function annotation of the UTI89 proteome.

### Functions enriched in uncharacterized proteins

To obtain a more comprehensive understanding of the biological functions of proteins encoded by *E. coli* UTI89, we used a log odds-ratio analysis (see the **Materials and Methods** section for details) to identify GO terms significantly enriched among poorly annotated proteins relative to well-annotated proteins in the UTI89 proteome (**Figures 5A**-**C**). For the BP and CC aspects, we were able to identify GO terms enriched among the poorly annotated set, thus reflecting annotations (and particular, pathways and biological functionalities) that are systematically missed in the previously existing UniProt-derived dataset. We showed the BP and CC GO terms with the strongest depletion and strongest enrichment among poorly annotated proteins in the UTI89 genome in **Supplementary Figures S2** to **S3**. For the MF aspect, there were no GO terms systematically enriched among the poorly annotated proteins, likely indicating that gaps in existing annotations do not reflect gaps in UniProt coverage of enzyme families or specific molecular-level roles, but rather, overall pathways that are either properly covered or missed. It is notable that several MF terms are depleted among the poorly annotated set, including ion binding, nucleotide binding, and core enzymatic activities such as transferase, hydrolase, oxidoreductase, and lyase, likely reflecting the relative ease of recognizing proteins belonging to these highly important and abundant categories.

**Figure 5.**
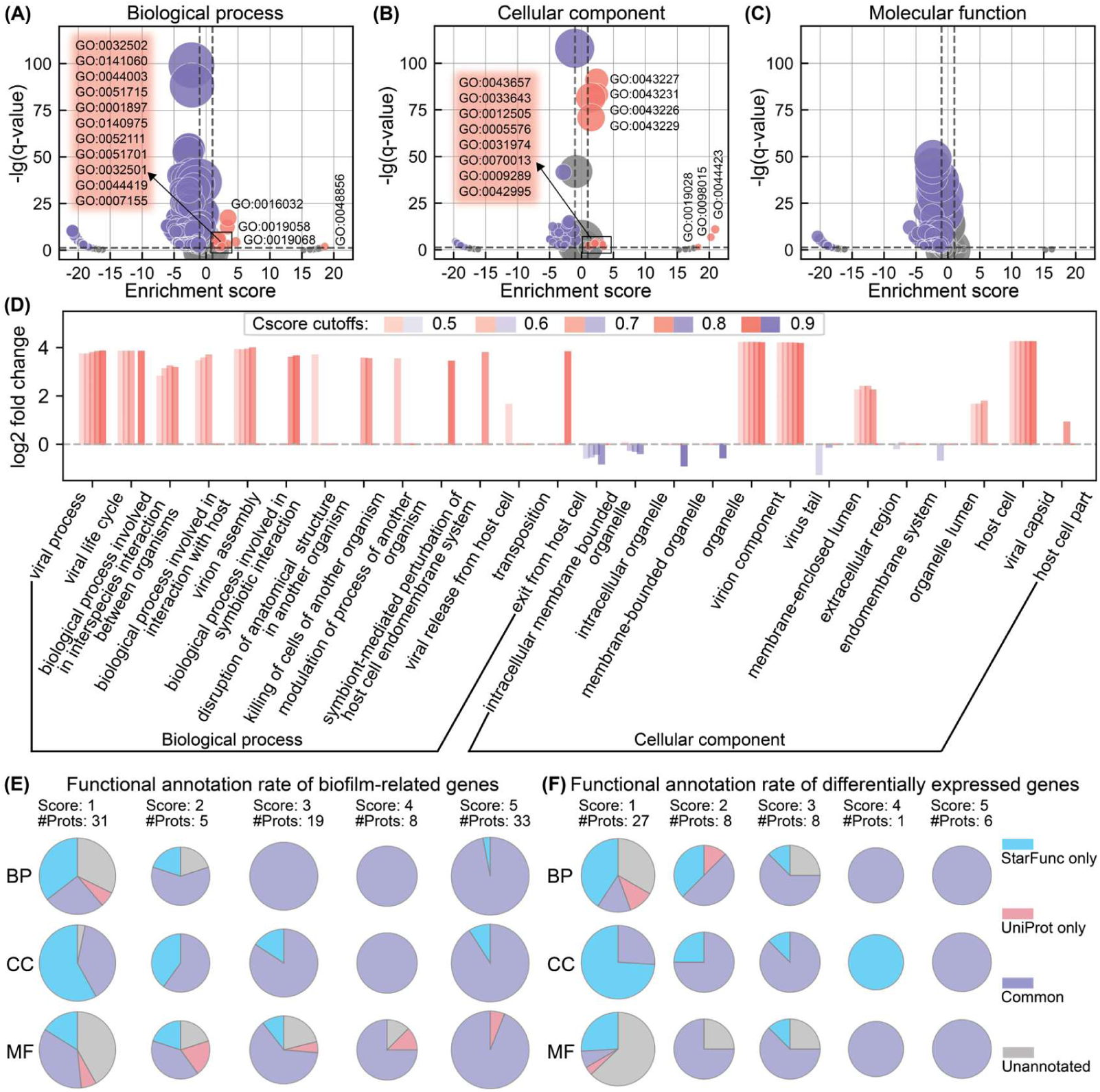
Enrichment of GO terms with a Cscore >0.5 across all three aspects in poorly- vs. well-annotated proteins and functional annotation rate. **(A)**, **(B)** and **(C)** are enrichment of GO terms for BP, CC, and MF, respectively. Each GO term is represented as a circle, with its *x*-position corresponding to the enrichment score among poorly annotated genes relative to well-annotated genes, its *y*-axis indicating the base-10 logarithm of the FDR-corrected *p*-value (*q*-value) for that enrichment, and the size proportional to the total number of genes in that GO term. GO terms with a *q*-value < 0.05 and an enrichment score > 1 are labeled with their corresponding GO term numbers. The descriptions and details for each GO term number can be found in **Supplementary Table S20**. **(D)** Assignment of GO terms enriched in poorly annotated proteins to phage-derived genes vs. other genes in the UTI89 genome. The *x*-axis shows GO terms (biological process and cellular component categories) that are present in phage-derived genes. Each bar within a GO term represents a different StarFunc Cscore cutoff. The *y*-axis indicates the log2 fold change, calculated as the number of phage-derived genes annotated with a given GO term divided by the number of genes in the entire UTI89 genome annotated with the same GO term, normalized by the proportion of phage-derived genes in the UTI89 genome (0.054). A positive value indicates enrichment of the GO term in phage-derived genes compared to the whole genome, while a negative value indicates depletion. **(E)** and **(F)** Functional annotation rates across annotation score categories for proteins involved in biofilm formation and proteins differentially expressed in biofilm-like intracellular bacterial communities (IBCs). Detailed numbers are provided in Supplementary **Tables S21** to **S26**.

### Enrichment of colonization- and virulence-associated BP GO terms

To explore the potential functional characteristics of poorly annotated proteins in UTI89 in terms of the biological process (BP), we performed GO term enrichment analysis on BP GO terms generated by StarFunc, as shown in **Figure 5A** and **Supplementary Figure S2**. We compared the enrichment patterns between well annotated and poorly annotated proteins by focusing on the GO terms significantly enriched at each StarFunc confidence threshold.

We observed that GO terms enriched in well-annotated proteins primarily involve fundamental and conserved cellular functions, such as “biosynthetic process/GO:0009058”, “macromolecule metabolic process/GO:0043170”, “gene expression/GO:0010467”, “RNA metabolic process/GO:0016070”, and “cellular component organization/GO:0016043”. These terms are frequently represented in well-characterized proteins and are consistent with core metabolic and housekeeping roles.

In contrast, GO terms enriched in poorly annotated proteins include “viral process/GO:0016032”, “viral life cycle/GO:0019058”, “virion assembly/GO:0019068”, “exit from host cell/GO:0035891”, and “viral release from host cell/GO:0019076”, suggesting that a large portion of these proteins are associated with prophage-derived sequences (the most likely origin for the abundance of virus-related GO terms). Additional terms such as “symbiont-mediated perturbation of host cell endomembrane system/GO:0052025” and “modulation of process of another organism/GO:0035821” point to functions probably involved in host-pathogen interactions. Similarly, we also identified GO terms related to pathogenic behavior and interspecies interactions, including “interaction with host/GO:0051701”, “interspecies interaction between organisms/GO:0044419”, “disruption of anatomical structure/GO:0141060”, “killing of cells of another organism/GO:0031640”, and “cell adhesion/GO:0007155”. These processes are indicative of bacterial strategies for colonizing and manipulating host environments; it is notable that in many such cases, the genes with these annotations also occur in prophages (**Figure 5D**), although it is in some cases unclear whether the assigned GO terms reflect a role in phage-bacteria or bacteria-host interactions (many cases of phage encoded genes involved in the latter interaction are well studied, including shiga toxin [49] and cholera toxin [50]). Among the genes annotated with the GO terms “interspecies interaction between organisms/GO:0044419” or “killing of cells of another organism/GO:0031640”, our pipeline also identified two *E. coli* encoded bacteria-host interaction genes: Q1R2T5 (hemolysin A), which is well-characterized, and Q1R2U0 (cytotoxic necrotizing factor 1), for which no BP annotation is currently available in UniProt despite its well characterized biological role in the relevant literature [51]. Similarly, the enrichment of proteins annotated with “cellular response to iron ion/GO:0071281” term further suggests deep involvement of poorly annotated genes in adaptation to host-imposed nutrient limitations [10], such as iron restriction during infection. Taken together, this indicates that poorly annotated proteins are not functionally random; instead, they are significantly enriched for biologically coherent GO terms, particularly those associated with phage-related and host-colonization processes; thus, our predictions reveal a wealth of new functionalities related both to the interface of UTI89 with the hosts that it parasitizes, and at the same time, with the viruses that parasitize it.

In addition, we observed an enrichment of new annotations for genes with two GO terms, “cell adhesion/GO:0007155” and “cellular response to iron ion/GO:0071281”, which arise entirely from non-prophage genes and play critical roles in UTI89 colonization, growth, and virulence [10], are enriched among poorly annotated proteins (see **Supplementary Figure S2**). Cell adhesion plays a critical role in UTI89 pathogenesis, enabling the bacteria to attach to uroepithelial cells, resist urinary flow [52], and establish intracellular communities [10, 15]. This function is also essential for biofilm formation, which provides protection against immune responses and antibiotic treatments, allowing UTI89 to establish persistent or recurrent infections [53]. Based on UniProt, a total of 61 UTI89 proteins are annotated with the GO term “cell adhesion”, comprising 26 well-annotated and 35 poorly annotated proteins. Beyond these, StarFunc predicted 8 additional proteins associated with this GO term, including 1 well-annotated and 7 poorly annotated proteins. Moreover, the consistent enrichment of “cell adhesion” across all Cscore thresholds (≥0.5 to ≥0.9) highlights the robustness of this result, providing new key protein candidates for investigating UTI89 colonization and survival. Another major aspect of host colonization by UPEC is overcoming nutrient deprivation-based host defense mechanisms. During infection, the host actively restricts the availability of iron ions through mechanisms involving transferrin and lipocalin 2 to inhibit bacterial growth [54]. As a UPEC strain, UTI89 activates iron-responsive mechanisms to counteract this nutritional stress [54]. The prediction of “cellular response to iron ion/GO: 0071281” in poorly annotated UTI89 proteins highlights their potential involvement in overcoming host-imposed iron limitation during infection. However, in UniProt, only 2 well annotated proteins with this GO term exist in the UTI89 proteome. StarFunc prediction provides other 8 proteins with this GO terms, and all of them come from poorly annotated proteins, providing potential novel targets involved in counteracting host defenses.

Taken together, our findings suggest that functional prediction for these proteins may reveal some biologically meaningful new patterns and underscores their likely importance in niche adaptation, virulence, and host colonization. At the same time, this also indicates, to some extent, the limitations of the current UniProt database in protein function annotation (a problem that is almost unavoidable, given the sheer volume of information that UniProt must aim to contain). These uncharacterized proteins may, in fact, serve as key drivers of phenotypic diversity and ecological versatility in pathogenic bacteria, and while in many cases the new StarFunc predictions are consistent with information that already exists somewhere in the literature, the advantage of our prediction sets is the automated provision of high-quality automated annotations (and associated structures) to supplement the information in existing databases.

### Enrichment of colonization- and virulence-associated CC GO terms

To explore the potential functional characteristics of poorly annotated proteins in UTI89 in terms of the cellular component (CC) aspect, we performed GO term enrichment analysis on CC GO terms generated by StarFunc under a series of Cscore cutoffs (0.5 to 0.9), as shown in **Figure 5B** and **Supplementary Figure S3**.

For well-annotated proteins, the enriched CC GO terms primarily correspond to highly conserved and well-characterized cellular structures and complexes. These included ribosomal components, membrane-associated complexes, enzyme-related complexes, and intracellular structural sites, all of which are core constituents of cellular machinery.

In contrast, GO terms enriched in poorly annotated proteins showed a different pattern. These proteins were associated with terms related to viral structures and secreted proteins. Enriched viral components included “virion component/GO:0044423”, “viral capsid/GO:0019028”, “virus tail/GO:0098015”, and host interaction terms such as “host cell/GO:0043657” and “host cell part/GO:0033643”. These GO terms are particularly enriched in phage-derived proteins, as shown in the CC part of **Figure 5D**. Moreover, we observed enrichment of terms associated with extracellular and membrane-bound activities, including “extracellular region/GO:0005576”, “pilus/GO:0009289”, “endomembrane system/GO:0012505”, “cell projection/GO:0042995”, and “membrane-enclosed lumen/GO:0031974”, many of which are likely involved in interactions at the host-pathogen interface. Additionally, we found that GO terms associated with intracellular and membrane-bounded organelles are predominantly enriched in bacteria-derived proteins as shown in the CC part of **Figure 5D** and may well represent secreted proteins that act in host cells.

Additionally, as shown in **Figure 5B** and **Supplementary Figure S3**, three virus-associated GO terms, “virion component/GO:0044423”, “virus tail/GO:0098015”, and “viral capsid/GO:0019028”, are consistently and significantly enriched in poorly annotated proteins across all Cscore cutoffs (from ≥0.5 to ≥0.9). This pattern is consistent with the results observed in the BP GO terms, suggesting that several prophages in the UTI89 genome are severely under-annotated. However, for the three GO terms, “virion component”, “virus tail”, and “viral capsid”, no UTI89 proteins are annotated with any of these terms in UniProt. In contrast, our pipeline predicted 51, 32, and 8 proteins with these GO terms, respectively. For two GO terms associated with bacteria-derived proteins, “pilus/GO:0009289” and “cell projection/GO:0042995”, significant enrichment was also observed in poorly annotated proteins across all Cscore cutoffs. The proteins associated with these two GO terms may play crucial roles in UTI89’s ability to adhere to host tissues, establish persistent colonization, and evade clearance from the urinary tract, thereby contributing significantly to its pathogenicity. In addition to the GO annotations provided by UniProt, our pipeline identified 1 additional protein associated with the pilus term and 17 with the cell projection term.

Together, these findings highlight a clear functional divergence between well-annotated and poorly annotated proteins. While the former are primarily linked to fundamental cellular processes, the latter appear to be enriched for proteins associated with extracellular functions, host interactions, and phage components. These patterns suggest that many uncharacterized proteins in UTI89 may be involved in virulence-associated or phage-related functions, thereby providing new candidate targets for future studies on bacterial pathogenicity and host-pathogen interactions.

The UPEC pathogenic cascade starts when bacteria in the lumen bind to bladder superficial umbrella cells using type 1 pili, which bind to mannosylated residues on host glycoproteins, specifically the mannosylated uroplakin Ia [55]. After binding, UPEC are able to invade the umbrella cells and establish intracellular bacterial communities (IBCs), which are biofilm-like communities that form in the first 6-12 hours post invasion [56]. IBCs are an essential step in virulence that enables evasion from host immune responses, and factors that promote biofilm development are associated with IBC formation [56]. To provide more detailed insights into the additional information revealed by our pipeline in the context of uropathogenesis, we checked functional annotation rates for two subsets of genes: 96 protein-coding genes associated with biofilm formation [57] and 50 differentially expressed genes identified in IBCs [58] (**Figures 5E** and **5F**). For the 96 biofilm-related genes, 36 had an annotation score of 1 or 2 in Uniprot; of this subset, our pipeline permitted the addition of high-confidence GO terms (Cscore >0.5) to all of them (30 new annotations for MF, 140 for BP and 133 for CC). With increasing annotation scores (4-5), most genes were already annotated, but StarFunc continued to provide additional functional terms even at score 5. These findings indicate that biofilm formation-related genes remain insufficiently annotated in existing databases, especially at low annotation scores, and that StarFunc substantially improves the number of proteins with specific GO terms. For the 50 genes identified as differentially expressed during IBC formation, 35 had an annotation score of 1 or 2 in UniProt. Within this subset, our pipeline enabled the addition of high-confidence GO term annotations (Cscore > 0.5) for 34 genes, yielding 81 new annotations in MF, 79 in BP, and 136 in CC. Notably, in the CC domain, StarFunc provided at least one specific functional annotation for nearly all poorly annotated proteins. Overall, these differentially expressed genes remain sparsely annotated, but StarFunc offers critical complementary information, underscoring its utility for uncovering functions of biofilm-associated, differentially expressed proteins.

### Case studies using the sequence-structure-function-derived function annotations

To provide examples of the actionable annotation information generated by our pipeline, we chose two representative case studies illustrating the integration of newly-derived predictions from our pipelines with previous knowledge.

#### *Case study 1: UTI89_C0931* (Q1RDZ8) is likely a phage lysis protein

Due to the potential biological relevance, and enrichment among poorly annotated genes, we focused on proteins with known or predicted annotations of “killing of cells of another organism/GO:0031640”, and thus compiled statistics based on annotations from UniProt and StarFunc. In our complete (post-pipeline) annotation set, a total of 21 proteins annotated with this GO term, with 13 out of them (61.9%) annotated by StarFunc and UniProt, 3 by UniProt only (14.3%), and 5 by StarFunc only (23.8%). All five proteins annotated exclusively by StarFunc had a UniProt annotation score of 1, and thus are essentially uncharacterized according to existing databases. The five genes encoding these proteins are in three distinct operons, and their positions in the UTI89 chromosome are illustrated in **Supplementary Figure S4**. From this set, we selected Q1RDZ8 as a case study, as it is an uncharacterized protein with minimal functional information available in UniProt (where it is annotated as a “DUF2570 domain-containing protein”, but has only a single annotated GO term, “membrane/GO:0016020” [and, by implication, its parent terms]). Even when using a more relaxed protein sequence identity threshold of 0.50 to identify better-annotated homologs, there are still no available matches in UniProt with more detailed annotations. In contrast, StarFunc provides high-confidence predictions for several GO terms such as “killing of cells of another organism/GO:0031640” and the more specific “symbiont-mediated cytolysis of host cell/GO:0001897”. Due to the absence of available templates in the PDB and the lack of high-pLDDT structure models in the AlphaFold DB, D-I-TASSER was used to predict the protein structure, which showed four α-helical regions connected by loops and arranged in an extended architecture (**Figure 6A**). When we used StoPred [59], a predictor for predicting the stoichiometry of the protein complex, to predict the possible oligomeric states (stoichiometries) of this protein, the top five predictions suggested that it may form dimeric, trimeric, tetrameric, 18-mer, and 10-mer assemblies (see **Figure 6B**). The corresponding AlphaFold3-predicted structures of these assemblies are shown in **Figure 6C**. Based on the AlphaFold3 results, the tetramer appears to be the most likely assembly form of Q1RDZ8, as it exhibits the highest ipTM value (0.29). Regardless of the stoichiometry, all predicted assemblies adopt rod-like coiled-coil architectures, which closely resemble the λ-phage Rz spanin system [60]. In this system, after degradation of the peptidoglycan layer by endolysin, the spanins are released, oligomerize (accumulation), and undergo conformational rearrangements that draw the inner and outer membranes into close proximity, leading to their fusion. This membrane fusion removes the outer membrane barrier, resulting in host cell lysis and phage release [60]. Computational analyses suggest that Q1RDZ8 adopts a spanin-like architecture and may mediate a similar membrane-fusion process, supported by StarFunc, StoPred, and AlphaFold3 predictions. Highly consistent with these findings, the operon containing Q1RDZ8 also encodes endolysin, holin, tail, and head completion/stabilization proteins. Collectively, these findings support that Q1RDZ8 is part of a phage lytic operon and has structural similarity to the λ phage spanin, although it had not been previously annotated as such.

**Figure 6.**
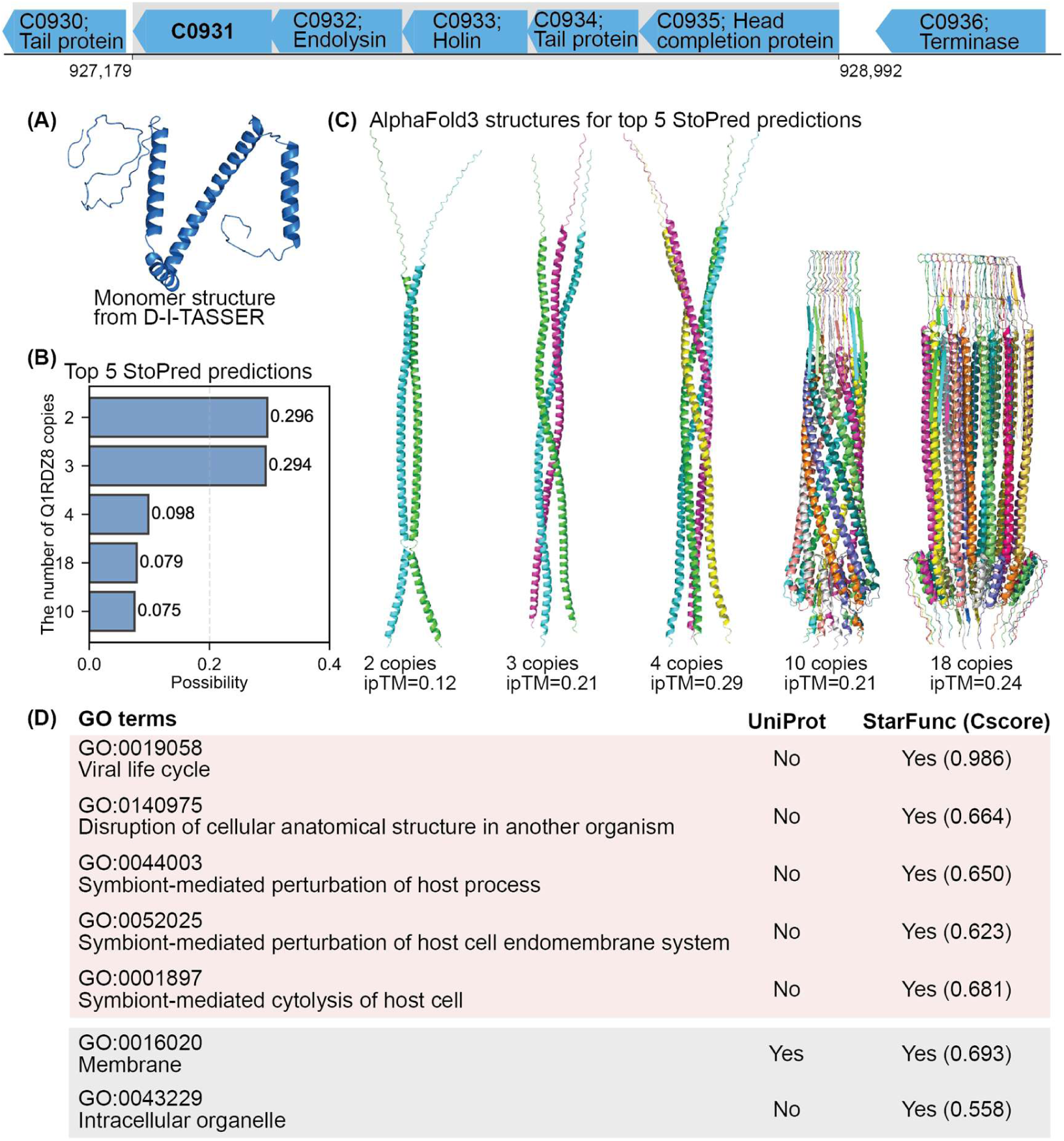
Predicted structure, stoichiometry and functions of the protein encoded by C0931 (Q1RDZ8). **(A)** Following our automated workflow, the monomer structure was modeled using D-I-TASSER. **(B)** Results of stoichiometry predictions on C0931 using StoPred. **(C)** AlphaFold3 structure predictions and accompanying confidence scores for the highest-likelihood StoPred stoichiometries. **(D)** The table lists the GO terms predicted by StarFunc, where pink and gray shading represent BP and CC, respectively, and only the leaf GO terms are shown (i.e., those without any child terms, thus representing the most specific functional predictions).

#### *Case study 2: ybtS* (Q1RAG5) is a critical virulence factor involved in siderophore biosynthesis

In the enrichment analysis related to BP terms, we also observed a significant enrichment of functions associated with the cellular response to iron ion (GO:0071281) among poorly annotated proteins. This finding is crucial for understanding the survival strategies of UTI89 in iron-restricted environments, including the bladder during infection. A total of 10 UTI89 proteins were found to be annotated with this GO term in our dataset. Among these, only one protein was annotated by both StarFunc and UniProt, one was exclusively annotated by UniProt, and eight solely by StarFunc. Among the eight proteins exclusively annotated by StarFunc, six were assigned a UniProt annotation score of 1, and two had a score of 2, thus reflecting assignment of the “response to iron ion” GO term to many poorly annotated proteins. The eight genes encoding these proteins are in five different operons (four on the UTI89 chromosome and one on the virulence plasmid); their positions in the UTI89 chromosome are illustrated in **Supplementary Figure S5**. We further investigated Q1RAG5 (UTI89_C2178) as a case study for the application of the predictions. Owing to the identification of a suitable homologous template from the PDB by DIAMOND [32, 33] in our automated pipeline workflow, the final structure was generated with LOMETS3 [22]. Although LOMETS3 does not provide confidence scores for its predicted protein structures, high-quality experimental structures in the PDB can serve as available templates for this protein. Consequently, we consider the structural model of Q1RAG5 generated by this method reliable. Notably, the sequence identity between Q1RAG5 and the template 5JY9 (chain A; a salicylate synthase from *Yersinia enterocloitica*) is above 99%. Based on the predicted structure and GO terms with high Cscores produced by StarFunc (**Figure 7**), we find that there is a clear discrepancy in the functional predictions provided by UniProt vs. StarFunc. Here, UniProt indicates the presence of anthranilate synthase activity (molecular function) and involvement in L-tryptophan biosynthesis (biological process), whereas StarFunc instead has several GO terms suggesting a role in siderophore biosynthesis and a role in iron homeostasis. The *ybtS* gene is part of the high pathogenicity island that encodes the genetic machinery necessary to make the siderophore yersiniabactin (Ybt), which was first identified in *Yersinia* species in the 1980s [61]. Ybt is a critical virulence factor that enables UPEC persistence within the nutrient-limited bladder environment, where the host sequesters iron as a form of nutritional immunity [62]. Ybt can bind to ferric iron, as well as other metals [63], in the external environment and then the Ybt-ferric iron complex is bound and imported by transporters in the outer membrane, where the metals are released by Ybt and available for bacterial use [63]. The *ybtS* gene product is responsible for the conversion of chorismic acid to salicylate in the cytoplasm, which is the first committed step of Ybt synthesis [61, 64]. The remaining steps of Ybt synthesis are performed by the orchestrated actions of six other genes in the high pathogenicity island [65]. Transcription of *ybtS*, as with the rest of the Ybt synthesis machinery, is regulated in response to metal and is induced in low-iron environments, as well as low concentrations of other metals like copper [63], consistent with the “cellular response to iron ion (GO:0071281) annotation arising from StarFunc (but absent in UniProt). Providing further consistency with this annotation, we found that supplementation with 2-hydroxybenzoate (salicylate), a downstream product of isochorismate synthase, restores yersiniabactin activity (**Figure 7C**). Furthermore, isotopic labeling experiments demonstrate that the pattern of labeling arising from deuterated salicylate in UTI89 Δ*ybtS* cells is consistent with direct utilization of the exogenous substrate in yersiniabactin production. (**Fig. 7D**).

**Figure 7.**
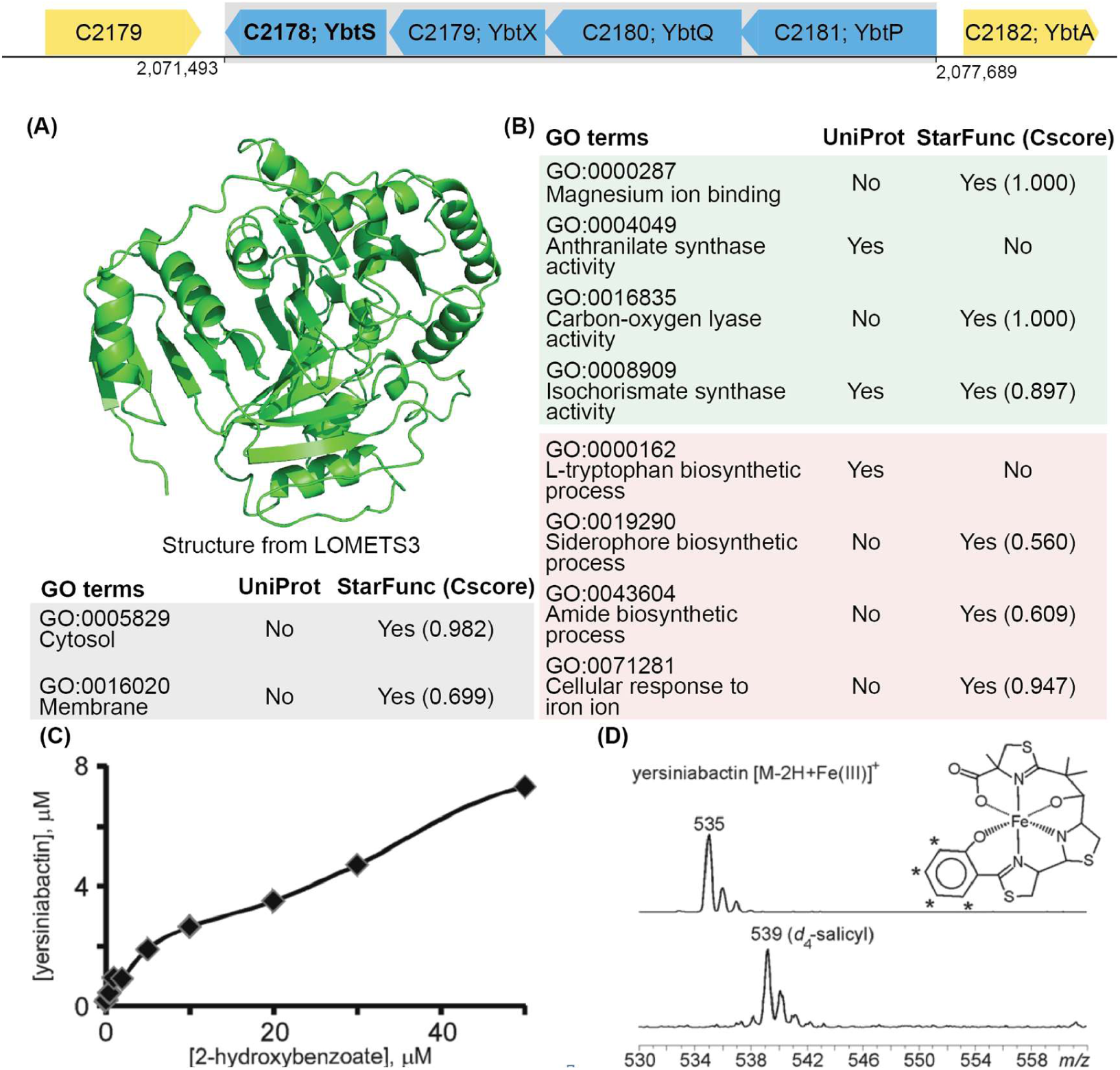
Predicted structure and functions of the protein encoded by Q1RAG5 (C2178). **(A)** Following our automated workflow, the structure was modeled using LOMETS3. **(B)** The leaf GO term predictions. **(C)** Yersiniabactin production is restored in UTI89Δ*ybtS* with chemical complementation by 2-hydroxybenzoic acid. **(D)** Chemical complementation with deuterated salicylic acid results in production of a yersiniabactin peak at m/z 539, consistent with incorporation of four nonexchangeable deuterium atoms at the designated positions (*), consistent with use of exogenous salicylic acid as a substrate for yersiniabactin biosynthesis.

The observed concordance between our predictions, literature evidence, and experimental validation reaffirms the efficacy and utility of our pipeline, and suggests that several of the extant GO annotations in UniProt are in error for this particular protein (as it is mis-assigned to a role in tryptophan biosynthesis instead of siderophore biosynthesis). Especially for poorly annotated proteins, our pipeline offers valuable insights, underscoring its significance in advancing our understanding of under-characterized genes, and especially in providing higher-quality automated annotations for use in high-throughput analysis.

### *E. coli* UTI89 structure-function database

We made full structure and function predictions for all 5,192 *E. coli* UTI89 proteins, available at https://seq2fun.dcmb.med.umich.edu/UTI89, as a user-friendly, searchable online database. On the main webpage, we summarize and list key information for each protein, including UniProt entry ID, protein name, gene name, protein size, annotation score, structure prediction method, and the GO terms, to facilitate quick lookup and filtering of target proteins, as shown in **Supplementary Figure S7**. We also provide detailed information for each target protein. On the individual protein pages, we show the top 20 structural analogs (10 from PDB and 10 from AFDB) aligned using Foldseek [66] and TM-align [67–69], the top 20 sequence homologs in UniProt aligned using BLASTp [70], and the StarFunc GO term prediction results, as shown in **Supplementary Figure S8**.

## Discussion

Here, we report the development and pilot application of an automated structure-based protein functional annotation pipeline as a solution to the growing inability of manual and experimental annotations to keep pace with the rapidly expanding space of known proteins. In this study, we developed and applied a cutting-edge pipeline for predicting protein structure and function across the entire *E. coli* proteome, encompassing 5,192 proteins. In addition, we established the UTI89 structure-function database, which, to the best of our knowledge, represents the first online resource to offer systematic function annotations based on proteome-wide structure modeling. For proteins lacking high-quality templates or models in PDB or AFDB, we employed the recently developed high-performance structure prediction methods, DMFold and D-I-TASSER [31], to generate new models. As a result, our pipeline improved pLDDT scores for 89% of the challenging subset of 696 proteins requiring *de novo* modeling, relative to AFDB models. For 13 proteins absent of structural information in UniProt, we provided DMFold-predicted structures with an average pLDDT of 82.79. Simultaneously, our pipeline can offer more detailed and specific functional annotations compared to those provided by UniProt. Moreover, we have found that the GO terms predicted by our pipeline can effectively complement the GO terms available from UniProt. Integrating both sets of annotations can further enhance the annotation coverage of the UTI89 proteome. Enrichment analysis of the predicted GO terms revealed that many poorly annotated proteins are involved in key processes related to microbial pathogenesis and host interaction. These include functions associated with adhesion, invasion, and acquisition of limiting nutrients such as iron, features that are central to understanding bacterial virulence and the dynamics of host-pathogen interactions. Our findings highlight the biological relevance of these uncharacterized proteins and the potential of our pipeline to uncover functionally critical components that are currently underrepresented in existing annotations. The comprehensive annotations of such proteins provided by our pipeline/database represent a crucial first step in identifying new potential disease biomarkers and even therapeutic targets. By analyzing two cases, *UTI89_C0931* and *ybtS*, we further demonstrated the performance and utility of our pipeline. Although our predictions have not been validated by new experiments, in both case studies existing literature substantively supports the predictions emerging from our pipelines, highlighting the extent to which our annotation pipeline can provide high-quality new annotations suitable for high-throughput analysis.

We anticipate that the database of structure/function predictions for the *E. coli* UTI89 proteome that we have built will serve as a significant resource for researchers. In addition, we are providing the fully open-source toolkit used for implementing these proteome-wide structure/function predictions, which will permit users to apply the same approach seamlessly to other organisms of interest. Nevertheless, we must note that we cannot assume that the predictions contained within this database are infallible, even though the performance of StarFunc meets or exceeds that of many state-of-the-art methods; careful consideration of the totality of available information is vital especially for informing follow-up experiments that might arise from a screen. In addition, the approach taken here will likely need to be updated frequently in order to make use of the strongest available set of tools for each of the involved subtasks. We have also not yet automated the pruning of some types of GO terms, such as those that are highly abundant, or those that are likely incorrect based on taxonomic analysis [38]; these and similar approaches might provide powerful future extensions of our workflow.

## Data and code availability

All data underlying this work is freely available at https://seq2fun.dcmb.med.umich.edu/UTI89 and the source code of the sequence-structure-function pipeline is freely available at https://github.com/QuanEvans/UTI89-Structure-Function-Pipeline.

## Acknowledgements

This work used the Advanced Cyberinfrastructure Coordination Ecosystem: Services and Support (ACCESS) program, which is supported by National Science Foundation [2138259, 2138286, 2138307, 2137603, and 2138296].

