## Supplementary Material for "Whole-proteome structure/function prediction in Uropathogenic *E. coli* reveals previously missed host-microbe and microbe-phage interaction pathways"

**Supplementary Texts**

- **Text S1**: Details of the construction of **Dataset I** and **Dataset II**.
- **Text S2**: Details of homology-based function transfer approach used in this study.
- **Text S3**: Definition of the Accretion-Weighted F-Measure (wFmax).

**Supplementary Tables**

**Table S1**: The details of 17 excluded common GO terms.
**Table S2**: The details of the obviously incorrect for *E. coli* UTI89.
**Table S3:** Primer and test sequences used to construct UTI89 mutants.
**Table S4**: Bacterial strains used in this study.
**Table S5**: Phage-related genes identified by PHASTER and filtered using bedtools with ≥90% genomic overlap.
**Table S6**: Functional annotation rates of whole UTI89 genome across annotation score categories for biological process.
**Table S7**: Functional annotation rates of whole UTI89 genome across annotation score categories for cellular component.
**Table S8**: Functional annotation rates of whole UTI89 genome across annotation score categories for molecular function.
**Table S9**: Functional annotation rates of UTI89 plasmid-encoded proteins across annotation score categories for biological process.
**Table S10**: Functional annotation rates of UTI89 plasmid-encoded proteins across annotation score categories for cellular component.
**Table S11**: Functional annotation rates of UTI89 plasmid-encoded proteins across annotation score categories for molecular function.
**Table S12**: Functional annotation rates of UTI89 proteins excluding phage-derived proteins across annotation score categories for biological process.
**Table S13**: Functional annotation rates of UTI89 proteins excluding phage-derived proteins across annotation score categories for cellular component.
**Table S14**: Functional annotation rates of UTI89 proteins excluding phage-derived proteins across annotation score categories for molecular function.
**Table S15**: Functional annotation rates of UTI89 phage-derived proteins across annotation score categories for biological process.
**Table S16**: Functional annotation rates of UTI89 phage-derived proteins across annotation score categories for cellular component.
**Table S17**: Functional annotation rates of UTI89 phage-derived proteins across annotation score categories for molecular function.
**Table S18**: pLDDT scores of the 5 proteins predicted by DMFold that also have available structure models in the AlphaFold Database (AFDB).
**Table S19**: pLDDT scores of the 13 proteins predicted by DMFold that haven't available structure models in the AFDB.
**Table S20**: The descriptions and details for each GO term enriched in poorly annotated proteins.
**Table S21**: Functional annotation rates of UTI89 biofilm formation-related proteins across annotation score categories for biological process.
**Table S22**: Functional annotation rates of UTI89 biofilm formation-related proteins across annotation score categories for cellular component.
**Table S23**: Functional annotation rates of UTI89 biofilm formation-related proteins across annotation score categories molecular function.

**Table S24**: Functional annotation rates of UTI89 proteins differentially expressed in biofilm-like intracellular bacterial communities (IBCs) across annotation score categories for biological process.
**Table S25**: Functional annotation rates of UTI89 proteins differentially expressed in biofilm-like intracellular bacterial communities (IBCs) across annotation score categories for cellular component.
**Table S26**: Functional annotation rates of UTI89 proteins differentially expressed in biofilm-like intracellular bacterial communities (IBCs) across annotation score categories for molecular function.

**Supplementary Figures**

**Figure S1**: Effect of models with different pLDDT scores on StarFunc-structure component GO term prediction performance.

**Figure S2**: Systematic enrichment and depletion of BP terms among poorly annotated *E. coli* UTI89 proteins.

**Figure S3**: Systematic enrichment and depletion of CC terms among poorly annotated *E. coli* UTI89 proteins.

**Figure S4**: Locations of the 7 operons on the UTI89 chromosome and the count of GO terms annotated for each operon by UniProt and StarFunc.

**Figure S5**: Locations of the 5 operons on the UTI89 chromosome and plasmid and the count of GO terms annotated for each operon by UniProt and StarFunc.

**Figure S6**: Liquid chromatography-mass spectrometry (LC-MS) reveals yersiniabactin as a major component of pellicle-conditioned media.

**Figure S7**: The main webpage of UTI89 structure-function database.

**Figure S8**: The details for each target protein in structure-function database.

#### Supplementary Texts

**Text S1**: Details of the construction of **Dataset I** and **Dataset II**.

The construction process of **Dataset I** involved three steps: (1) collecting 9,977 proteins that received new GO annotations in the UniProt-GOA 2025-05 release but had no annotations in the same GO aspect in the 2024-04 release; (2) removing redundancy using CD-HIT at a 40% sequence identity threshold and 80% alignment coverage; and (3) fifty proteins were randomly selected from each group of targets determined by AlphaFold2, Modeller, LOMETS, D-I-TASSER, and DMFold according to the workflow, resulting in the final test set with 250 proteins.

The construction process of **Dataset II** involved: (1) based on the data processing and selection procedures for **Dataset I**, 500 proteins were selected, including an additional 250 proteins; (2) for each of these proteins, 1,000 structure models were generated by running AlphaFold2 200 times with different random seeds, 100 runs with default settings and 100 runs without MSA or template information; (3) those proteins for which the 1,000 models were distributed across the three pLDDT intervals: (0, 50], (50, 70], and (70, 100] were selected as test proteins. In total, 128 proteins met these criteria and were selected as the final **Dataset II** to investigate the relationship between pLDDT and the performance of StarFunc.

**Text S2**: Details of homology-based function transfer approach used in this study

For sequence-based method, query protein sequence is searched against the UniProt Gene Ontology Annotation (UniProt-GOA) database by BLASTp [1] to identify homologous templates with function annotations, and protein functions of templates are transferred to query protein based on the bit-score and sequence identify of BLASTp hit. The prediction confidence score ($Cscore_{seq}$) for each GO term $q$ is calculated as follows,

$$Cscore_{\text{seq}}(q)=\frac{\sum_{k=1}^{K(q)} bitscore_k(q)\times ID_k(q)}{\sum_{k=1}^{K} bitscore_k\times ID_k} \quad \text{(S1)}$$

where $bitscore_k$ and $ID_k$ are the bit-score and sequence identity of the $k$-th BLASTp hit, while $K$ is the total number of hits with at least one GO term in the same GO aspect (Molecular Function, MF; Biological Process, BP; or Cellular Component, CC) as term $q$. Meanwhile, $bitscore_k(q)$, $ID_k(q)$ and $K(q)$ are the corresponding values for templates with GO term $q$. Here, the sequence identity ($ID_k$) for hit $k$ is calculated as the equation S2:

$$ID_k=\frac{Liden_k}{\max(L,L_k)} \quad \text{(S2)}$$

Here, $L$ and $L_k$ are the length of the query and the $k$-th template sequence respectively, while $Liden_k$ is the number of identical amino acids between the two proteins in the BLASTp alignment. Based on our recent benchmark study

[2], combining both bit-score and sequence identity yields better performance than using either metric alone.

When generating the UniProt-GOA sequence database, we only use experimental GO annotations as defined by CAFA [3] (evidence codes: EXP, IDA, IPI, IMP, IGI, IEP, HTP, HDA, HMP, HGI, HEP, TAS, and IC), excluding computational prediction such as those with evidence code IEA. Negative annotations indicating lack of protein function (with qualifier “NOT” or evidence codes “ND”) are also excluded. Parent GO terms are automatically propagated from the leaf terms annotated by UniProtGOA. Similar to the practice in previous CAFA challenges [3], protein binding-only templates, i.e., template proteins whose only Molecular Function (MF) leaf term is GO:0005515 “protein binding”, are not used for MF term prediction, because “protein binding” is a highly generic function description.

**Text S3**: Definition of the Accretion-Weighted F-Measure ($wF_{max}$)

The metric $wF_{max}$ represents the maximum value of the protein-centric, information accretion-weighted F-measure across all prediction thresholds ($\tau$). It is defined as:

$$wF_{max} = \max_{\tau\in(0,1]}\left(\frac{2wpr(\tau)\times wrc(\tau)}{wpr(\tau)+wrc(\tau)}\right) \tag{S3}$$

$$wpr(\tau) = \frac{1}{m(\tau)}\sum_{i=1}^{m(\tau)}\frac{\sum_q IA(q)\times 1(q\in P_i(\tau)\wedge T_i)}{\sum_q IA(q)\times 1(q\in P_i(\tau))} \tag{S4}$$

$$wrc(\tau) = \frac{1}{n}\sum_{i=1}^{n_e}\frac{\sum_q IA(q)\times 1(q\in P_i(\tau)\wedge T_i)}{\sum_q IA(q)\times 1(q\in T_i)} \tag{S5}$$

$$IA(q)=\log_2\left(\frac{1+|\text{proteins with parent term(s) of } q|}{1+|\text{proteins with term } q|}\right) \tag{S6}$$

where $wpr(\tau)$ and $wrc(\tau)$ denote the information accretion-weighted precision and recall, respectively, at a given prediction score threshold $\tau$. $P_i(\tau)$ represents the set of predicted GO terms for protein $i$ with confidence scores equal to or greater than $\tau$, and $T_i$ denotes the set of ground-truth GO terms for protein $i$. The value $m(\tau)$ indicates the number of proteins with at least one predicted GO term scoring above $\tau$, while $n$ is the total number of proteins in the benchmark. 1 is the indicator function. $q$ denotes the $q$-th GO term, and $IA(q)$ refers to its information accretion [4].

**Text S5**: Ferric yersiniabactin uptake by the tonB-dependent outer membrane transporter does not mimic the UTI89Δ*ybtS* phenotype.

To determine whether iron scavenging relates yersiniabactin dependence to the normal pellicle phenotype, we examined pellicle formation by UTI89Δ*fyuA*, which lacks the tonB-dependent ferric yersiniabactin outer membrane transporter. As expected, growth of UTI89Δ*fyuA* in iron-chelated medium containing ferric yersiniabactin as the sole iron source was deficient when compared to wild type UTI89 or UTI89Δ*ybtS*. UTI89Δ*fyuA* did not, however, reproduce the yersiniabactin-deficient pellicle phenotype and was instead grossly similar to wild type UTI89. This finding suggests that the effect of yersiniabactin upon pellicle formation is attributable to yersiniabactin biosynthesis and not subsequent reuptake of its ferric complex.

#### Supplementary Tables

**Table S1.** The details of 17 common GO terms that were excluded due to their high prevalence.

| Aspects of GO terms | GO term numbers | Description |
|---|---|---|
| Biological process | GO:0008150 | Biological process |
| | GO:0009987 | Cellular process |
| | GO:0008152 | Metabolic process |
| | GO:0044237 | Cellular metabolic process |
| | GO:0044238 | Primary metabolic process |
| | GO:0050896 | Response to stimulus |
| Molecular function | GO:0003674 | Molecular function |
| | GO:0005488 | Binding |
| | GO:0005515 | Protein binding |
| | GO:0003824 | Catalytic activity |
| | GO:0036094 | Small molecule binding |
| | GO:0097159 | Organic cyclic compound binding |
| | GO:1901363 | Heterocyclic compound binding |
| | GO:0043167 | Ion binding |
| Cellular component | GO:0005575 | Cellular component |
| | GO:0110165 | Cellular anatomical entity |
| | GO:0005622 | Intracellular anatomical structure |

**Table S2.** The details of the obviously incorrect (for *E. coli*) GO terms excluded from further analysis.

| Aspects of GO terms | GO term numbers | Description |
|---|---|---|
| Molecular function | GO:0043886 | Structural constituent of carboxysome shell |
| Cellular component | GO:0005739 | Mitochondrion |
| | GO:0031981 | Nuclear lumen |
| | GO:0005634 | Nucleus |

**Table S3.** Primer and test sequences used to construct UTI89 mutants.

| name | position | sequence |
|---|---|---|
| fur | left | 5’-TCAGGCTGGCTTATTTGCCTTCGTGCGCGTGCTCAT |
| | | CTTCGCGGCAATCGGTGTAGGCTGGAGCTGCTTC-3’ |
| | right | 5’-AGTAACAGGACAGATTCCGCATGACTGATAACAAT |
| | | ACCGCCCTAAAGAAACATATGAATATCCTCCTTAG-3’ |
| fur test | left | 5’-AGGCAACGCAAACCGGAAAT-3’ |
| | right | 5’-CAAGTGGCCTTGCCGTTGTA-3’ |
| ybtE | left | 5’-ATGAATTCTTCCTTTGAATCTCTGATTGAACAGTAT |
| | | CCCTTACCCATTGCGTGTAGGCTGGAGCTGCTTC-3’ |
| | right | 5’-TATTTCAACCTGTTTCGGGTCGGTTTGCGCTTATT |
| | | GGGCAGAATGGCGATCATATGAATATCCTCCTTAG-3’ |
| ybtE test | left | 5’-ACGCAGATTGTCCGCCATTT-3’ |
| | right | 5’-CCCTGTAAGACGGCGAAACG-3’ |
| fyuA | left | 5’-TCTTACAGGGACTCACAACAATGAAAATGACACGG |
| | | CTTTATCCTCTGGCCCATATGAATATCCTCCTTAG-3’ |
| | right | 5’-TCAGAAGAAATCAATTCGCGTATTGATACCGACGGT |
| | | GCGACCCATATTGAGTGTAGGCTGGAGCTGCTTC-3’ |
| fyuA test | left | 5’-ACCGACCCGAAACAGGTTGA-3’ |
| | right | 5’-CCGAATGATTTTCGGGGTGA-3’ |

**Table S4.** Bacterial strains used in this study.

| Strain | Description | Reference |
|---|---|---|
| UTI89 | human cystitis isolate | ref Mulvey 2001 [5] |
| UTI89Δ*fur* | *fur* knockout | this study |
| UTI89Δ*entB* | *entB* knockout | ref Henderson 2009 [6] |
| UTI89Δ*iroB* | *entB* knockout | ref Henderson 2009 [6] |
| UTI89Δ*ybtS* | *ybtS* knockout | ref Henderson 2009 [6] |
| UTI89Δ*ybtE* | *ybtE* knockout | this study |
| UTI89Δ*yfuA* | *fyuA* knockout | this study |

**Table S6.** Functional annotation rates of whole UTI89 genome across annotation score categories for biological process.

| | Score 1 | Score 2 | Score 3 | Score 4 | Score 5 |
|---|---|---|---|---|---|
| StarFunc only | 39.3% | 25.2% | 10.3% | 6.2% | 1.8% |
| Common | 14.7% | 50.8% | 81.4% | 90.4% | 96.5% |
| UniProt only | 3.5% | 7.5% | 4.6% | 1.8% | 1.0% |
| Unannotated | 42.5% | 16.5% | 3.7% | 1.6% | 0.7% |

**Table S7.** Functional annotation rates of whole UTI89 genome across annotation score categories for cellular component.

| | Score 1 | Score 2 | Score 3 | Score 4 | Score 5 |
|---|---|---|---|---|---|
| StarFunc only | 75.2% | 33.1% | 24.6% | 15.0% | 9.2% |
| Common | 22.7% | 66.6% | 75.4% | 85.0% | 90.8% |
| UniProt only | 0.2% | - | - | - | - |
| Unannotated | 1.9% | 0.3% | - | - | - |

**Table S8.** Functional annotation rates of whole UTI89 genome across annotation score categories for molecular function.

| | Score 1 | Score 2 | Score 3 | Score 4 | Score 5 |
|---|---|---|---|---|---|
| StarFunc only | 20.4% | 17.1% | 7.1% | 2.8% | 0.3% |
| Common | 21.3% | 56.2% | 74.7% | 88.9% | 96.5% |
| UniProt only | 3.9% | 6.0% | 6.7% | 3.9% | 1.7% |
| Unannotated | 54.4% | 20.7% | 11.5% | 4.4% | 1.5% |

**Table S9.** Functional annotation rates of UTI89 plasmid-encoded proteins across annotation score categories for biological process.

| | Score 1 | Score 2 | Score 3 | Score 4 | Score 5 |
|---|---|---|---|---|---|
| StarFunc only | 32.6% | 30.0% | - | - | - |
| Common | 15.2% | 40.0% | 100.0% | - | - |
| UniProt only | 3.0% | 20.0% | - | - | - |
| Unannotated | 49.2% | 10.0% | - | - | - |

**Table S10.** Functional annotation rates of UTI89 plasmid-encoded proteins across annotation score categories for cellular component.

| | Score 1 | Score 2 | Score 3 | Score 4 | Score 5 |
|---|---|---|---|---|---|
| StarFunc only | 75.8% | 30.0% | - | - | - |
| Common | 22.7% | 70.7% | 100.0% | - | - |
| UniProt only | - | -% | - | - | - |
| Unannotated | 1.5% | -% | - | - | - |

**Table S11.** Functional annotation rates of UTI89 plasmid-encoded proteins across annotation score categories for molecular function.

| | Score 1 | Score 2 | Score 3 | Score 4 | Score 5 |
|---|---|---|---|---|---|
| StarFunc only | 22.0% | 10.0% | - | - | - |
| Common | 19.7% | 60.0% | 100.0% | - | - |
| UniProt only | 6.0% | 10.0% | - | - | - |
| Unannotated | 52.3% | 20.0% | - | - | - |

**Table S12.** Functional annotation rates of UTI89 proteins excluding phage-derived proteins across annotation score categories for biological process.

| | Score 1 | Score 2 | Score 3 | Score 4 | Score 5 |
|---|---|---|---|---|---|
| StarFunc only | 38.8% | 25.7% | 10.4% | 6.1% | 1.8% |
| Common | 14.2% | 50.4% | 81.3% | 90.5% | 96.5% |
| UniProt only | 3.6% | 7.0% | 4.6% | 1.8% | 0.9% |
| Unannotated | 43.4% | 16.9% | 3.7% | 1.6% | 0.8% |

**Table S13.** Functional annotation rates of UTI89 proteins excluding phage-derived proteins across annotation score categories for cellular component.

| | Score 1 | Score 2 | Score 3 | Score 4 | Score 5 |
|---|---|---|---|---|---|
| StarFunc only | 73.6% | 32.1% | 24.1% | 14.3% | 9.2% |
| Common | 25.1% | 67.8% | 75.9% | 85.7% | 90.8% |
| UniProt only | - | - | - | - | - |
| Unannotated | 1.3% | 0.1% | - | - | - |

**Table S14.** Functional annotation rates of UTI89 proteins excluding phage-derived proteins across annotation score categories for molecular function.

| | Score 1 | Score 2 | Score 3 | Score 4 | Score 5 |
|---|---|---|---|---|---|
| StarFunc only | 19.4% | 17.7% | 7.1% | 2.8% | 0.3% |
| Common | 21.5% | 55.1% | 74.5% | 88.8% | 96.5% |
| UniProt only | 3.6% | 5.9% | 6.8% | 4.0% | 1.7% |
| Unannotated | 55.5% | 21.3% | 11.6% | 4.4% | 1.5% |

**Table S15.** Functional annotation rates of UTI89 phage-derived proteins across annotation score categories for biological process.

| | Score 1 | Score 2 | Score 3 | Score 4 | Score 5 |
|---|---|---|---|---|---|
| StarFunc only | 42.8% | 13.1% | - | 20.0% | - |
| Common | 18.2% | 60.9% | 100.0% | 80.0% | 85.7% |
| UniProt only | 3.0% | 21.7% | - | - | 14.3% |
| Unannotated | 36.0% | 4.3% | - | - | - |

**Table S16.** Functional annotation rates of UTI89 phage-derived proteins across annotation score categories for cellular component.

| | Score 1 | Score 2 | Score 3 | Score 4 | Score 5 |
|---|---|---|---|---|---|
| StarFunc only | 86.5% | 60.9% | 66.7% | 100.0% | 14.3% |
| Common | 5.9% | 34.8% | 33.3% | - | 85.7% |
| UniProt only | 1.7% | - | - | - | - |
| Unannotated | 5.9% | 4.3% | - | - | - |

**Table S17.** Functional annotation rates of UTI89 phage-derived proteins across annotation score categories for molecular function.

| | Score 1 | Score 2 | Score 3 | Score 4 | Score 5 |
|---|---|---|---|---|---|
| StarFunc only | 27.1% | - | - | - | - |
| Common | 19.9% | 87.0% | 100.0% | 100.0% | 100.0% |
| UniProt only | 6.4% | 8.7% | - | - | - |
| Unannotated | 46.6% | 4.3% | - | - | - |

**Table S18.** pLDDT scores of the 5 proteins predicted by DMFold that also have available structure models in the AlphaFold Database (AFDB).

| UinProt ID | pLDDT | |
|---|---|---|
| | DMFold | AFDB |
| Q1RD73 | 58.93 | 62.31 |
| Q1R2T5 | 70.46 | 68.96 |
| Q1R1Q5 | 74.17 | 69.28 |
| Q1R286 | 73.86 | 50.22 |
| Q1RE11 | 67.98 | 56.90 |
| Average | 69.08 | 61.53 |
| Median | 70.46 | 62.31 |

**Table S19.** pLDDT scores of the 13 proteins predicted by DMFold that haven't available structure models in the AFDB.

| UinProt ID | pLDDT | |
|---|---|---|
| | DMFold | AFDB |
| Q1RAC4 | 79.19 | - |
| Q1R2R5 | 85.09 | - |
| Q1RAD0 | 83.29 | - |
| Q1RAD2 | 80.73 | - |
| Q1RAD3 | 82.77 | - |
| Q1RAF9 | 83.70 | - |
| Q1RE24 | 65.83 | - |
| Q1R4Z3 | 87.44 | - |
| Q1R728 | 82.46 | - |
| Q1R8M6 | 91.82 | - |
| Q1RAD6 | 83.51 | - |
| Q1RC09 | 88.05 | - |
| Q1RFP2 | 82.43 | - |
| Average | 82.79 | - |
| Median | 83.29 | - |

**Table S21.** Functional annotation rates of UTI89 biofilm formation-related proteins across annotation score categories for biological process.

| | Score 1 | Score 2 | Score 3 | Score 4 | Score 5 |
|---|---|---|---|---|---|
| StarFunc only | 35.5% | 20.0% | - | - | 3.0% |
| Common | 25.8% | 60.0% | 100.0% | 100.0% | 96.7% |
| UniProt only | 6.5% | - | - | - | - |
| Unannotated | 32.2% | 20.0% | - | - | - |

**Table S22.** Functional annotation rates of UTI89 biofilm formation-related proteins across annotation score categories for cellular component.

| | Score 1 | Score 2 | Score 3 | Score 4 | Score 5 |
|---|---|---|---|---|---|
| StarFunc only | 58.1% | 40.0% | 15.8% | - | 9.1% |
| Common | 38.7% | 60.0% | 84.2% | 100.0% | 90.9% |
| UniProt only | - | - | - | - | - |
| Unannotated | 3.2% | - | - | - | - |

**Table S23.** Functional annotation rates of UTI89 biofilm formation-related proteins across annotation score categories for molecular function.

| | Score 1 | Score 2 | Score 3 | Score 4 | Score 5 |
|---|---|---|---|---|---|
| StarFunc only | 16.1% | 20.0% | 10.5% | - | - |
| Common | 35.5% | 40.0% | 63.2% | 75.0% | 93.9% |
| UniProt only | 6.5% | 20.0% | 5.3% | 12.5% | 6.1% |
| Unannotated | 41.9% | 20.0% | 21.0% | 12.5% | - |

**Table S24.** Functional annotation rates of UTI89 proteins differentially expressed in biofilm-like intracellular bacterial communities (IBCs) across annotation score categories for biological process.

| | Score 1 | Score 2 | Score 3 | Score 4 | Score 5 |
|---|---|---|---|---|---|
| StarFunc only | 40.8% | 37.5% | 12.5% | - | - |
| Common | 14.8% | 50.0% | 62.5% | 100.0% | 100.0% |
| UniProt only | 11.1% | 12.5% | - | - | - |
| Unannotated | 33.3% | - | 25.0% | - | - |

**Table S25.** Functional annotation rates of UTI89 proteins differentially expressed in biofilm-like intracellular bacterial communities (IBCs) across annotation score categories for cellular component.

| | Score 1 | Score 2 | Score 3 | Score 4 | Score 5 |
|---|---|---|---|---|---|
| StarFunc only | 74.1% | 25.0% | 12.5% | 100.0% | - |
| Common | 25.9% | 75.0% | 87.5% | - | 100.0% |
| UniProt only | - | - | - | - | - |
| Unannotated | - | - | - | - | - |

**Table S26.** Functional annotation rates of UTI89 proteins differentially expressed in biofilm-like intracellular bacterial communities (IBCs) across annotation score categories for molecular function.

| | Score 1 | Score 2 | Score 3 | Score 4 | Score 5 |
|---|---|---|---|---|---|
| StarFunc only | 25.9% | - | 12.5% | - | - |
| Common | 7.4% | 75.0% | 62.5% | 100.0% | 100.0% |
| UniProt only | 3.7% | - | - | - | - |
| Unannotated | 63.0% | 25.0% | 25.0% | - | - |

### Supplementary Figures

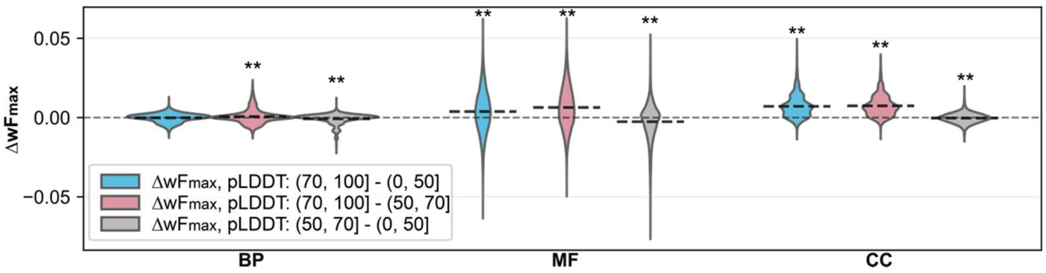


**Figure S1.** Effect of models with different pLDDT scores on StarFunc-structure component GO term prediction performance. Asterisks denote statistical significance levels: * indicates $p < 0.05$ and ** indicates $p < 0.01$, and dash line denotes the mean $\Delta wF_{max}$.

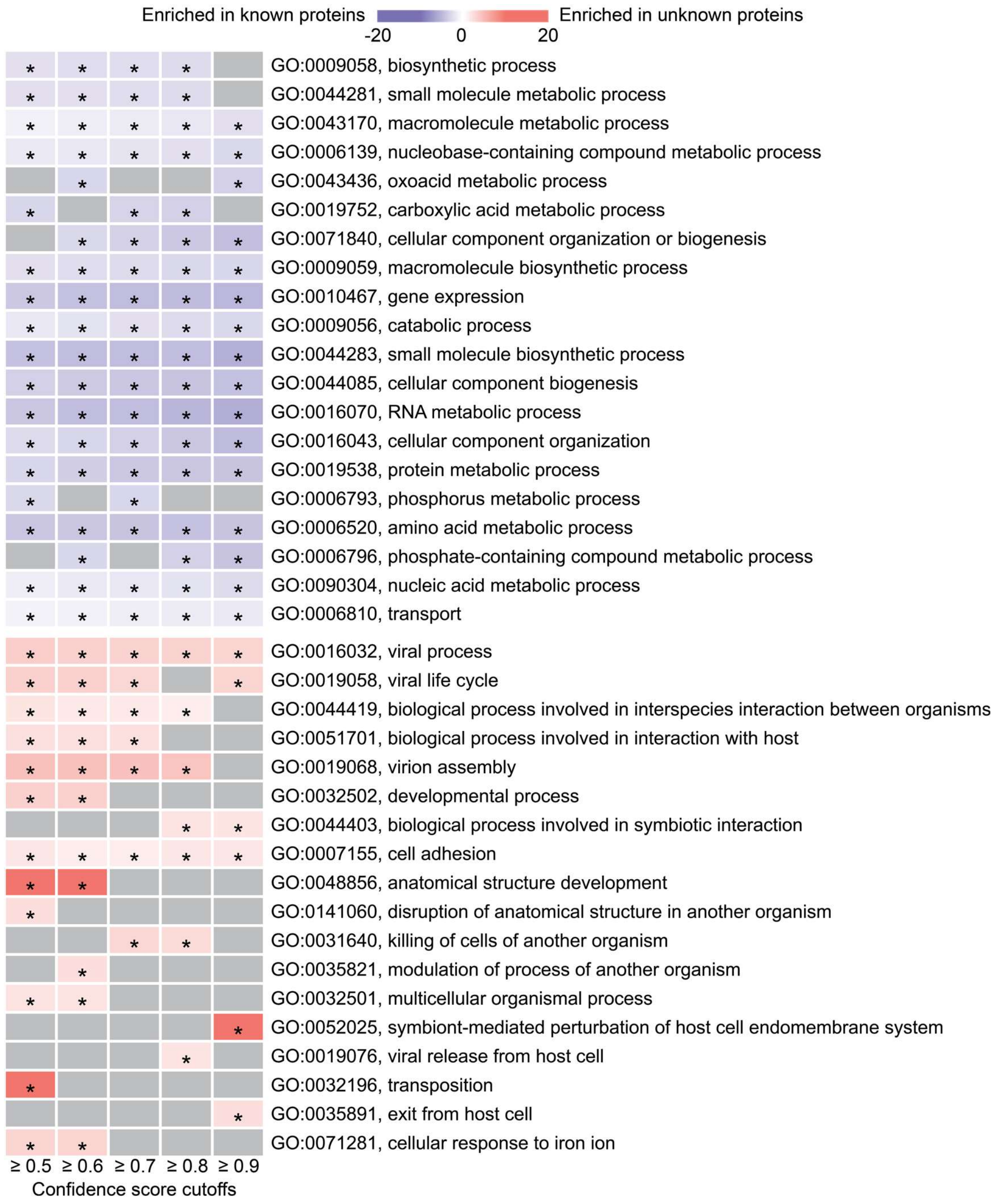


**Figure S2.** Systematic enrichment and depletion of BP terms among poorly annotated *E. coli* UTI89 proteins. The top and bottom panels display the most strongly under- and over-represented GO terms, respectively, ranked by *q*-value. Cells are colored according to the enrichment score for each GO term, and the asterisk represents the significance (*q*-value <0.05). The results are presented across five different Cscore cutoffs of StarFunc. Higher Cscores represent higher confidence for the predicted GO terms. Cells colored in grey without an asterisk indicate that the corresponding GO term does not reach statistical significance at the given Cscore cutoff.

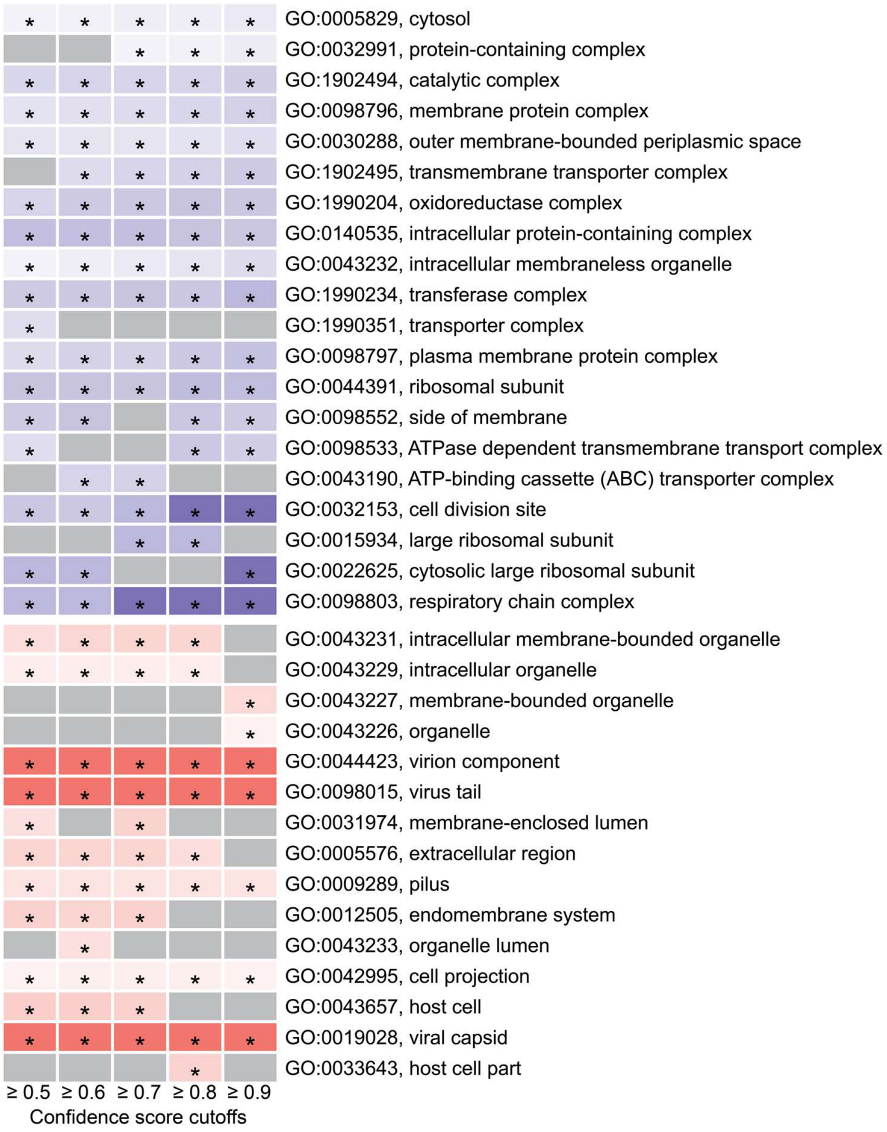


**Figure S3.** Systematic enrichment and depletion of CC terms among poorly annotated *E. coli* UTI89 proteins.

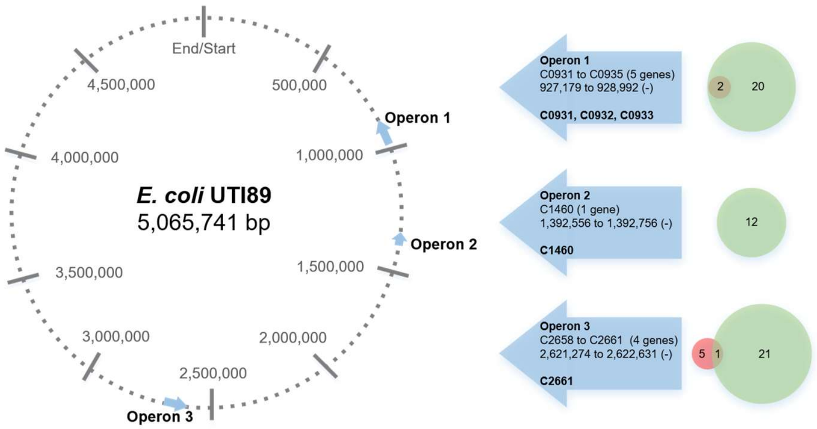


**Figure S4.** Locations of the 3 operons on the UTI89 chromosome with 'killing of cells of another organism/GO:0031640' annotations, and the count of GO terms annotated for each operon by UniProt and StarFunc. The first line in each arrow represents the order of the operon, the second line lists the genes it contains, the third specifies its precise location in the chromosome, and the fourth represents the 'target genes' within the operon. Here, 'target genes' refer to those exclusively annotated by StarFunc with 'killing of cells of another organism/GO:0031640'. Red circles represent the total number of GO terms provided by UniProt for each operon, green circles denote the total number provided by StarFunc, and the overlapping area shows the count of GO terms common to both.

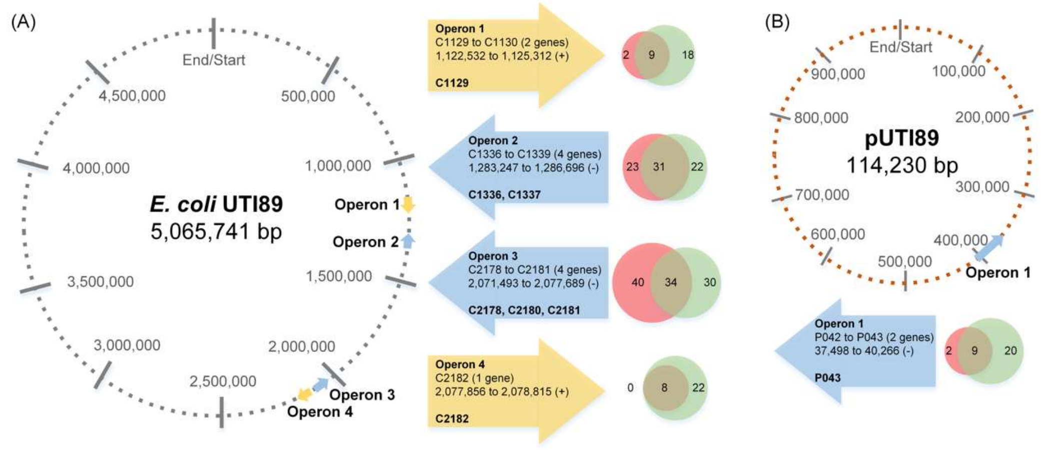


**Figure S5.** Locations of the 5 operons containing at least one gene with the "cellular response to iron iron/GO:0071281" GO term on the UTI89 chromosome and plasmid, and the count of GO terms annotated for each operon by UniProt and StarFunc. **(A)** Locations of the 4 operons on the UTI89 chromosome and **(B)** Locations of the 1 operon on the UTI89 plasmid DNA. Here, 'target genes' refer to those exclusively annotated by StarFunc with "cellular response to iron ion".

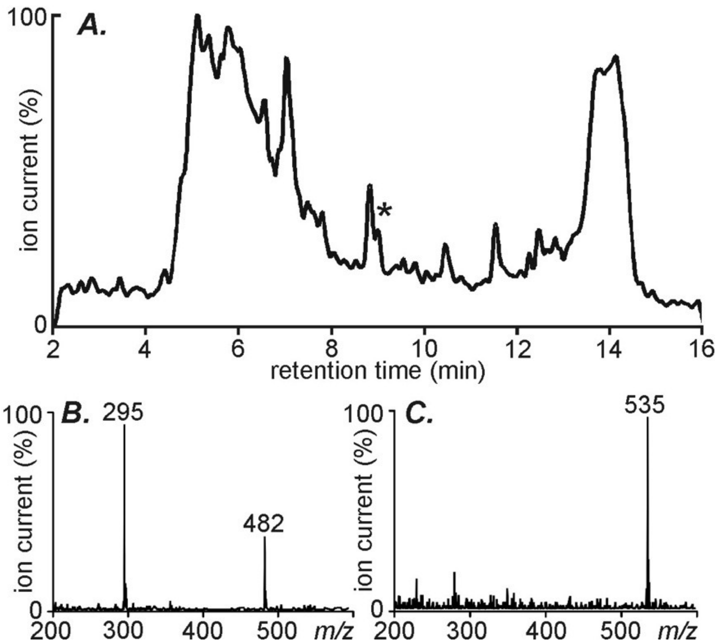


**Figure S6.** Liquid chromatography-mass spectrometry (LC-MS) reveals yersiniabactin as a major component of pellicle-conditioned media. We used LC-MS to determine what prominent metabolites are secreted by UTI89 pellicles. (A) Comparison to unconditioned media revealed that the pellicle culture produced a prominent peak at 9 minutes (indicated by *). (B) A full scan mass spectrum of this peak revealed major ions at *m/z* 482 and 295, consistent with the $[M+H]^+$ for yersiniabactin and its source decomposition product, respectively. (C) At 7.6 minutes, the corresponding ferric yersiniabactin ion was also observed at *m/z* 535.

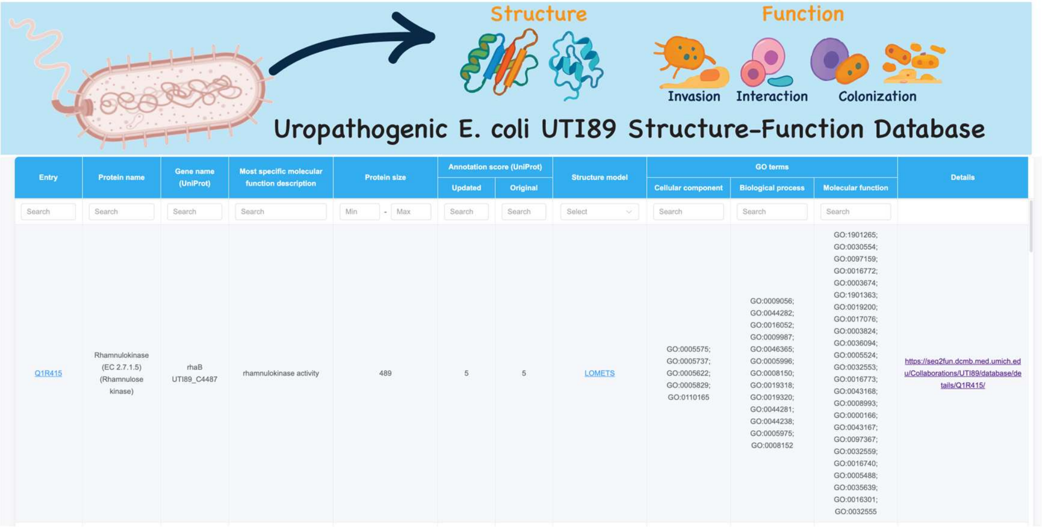


**Figure S7.** The main webpage of UTI89 structure-function database.

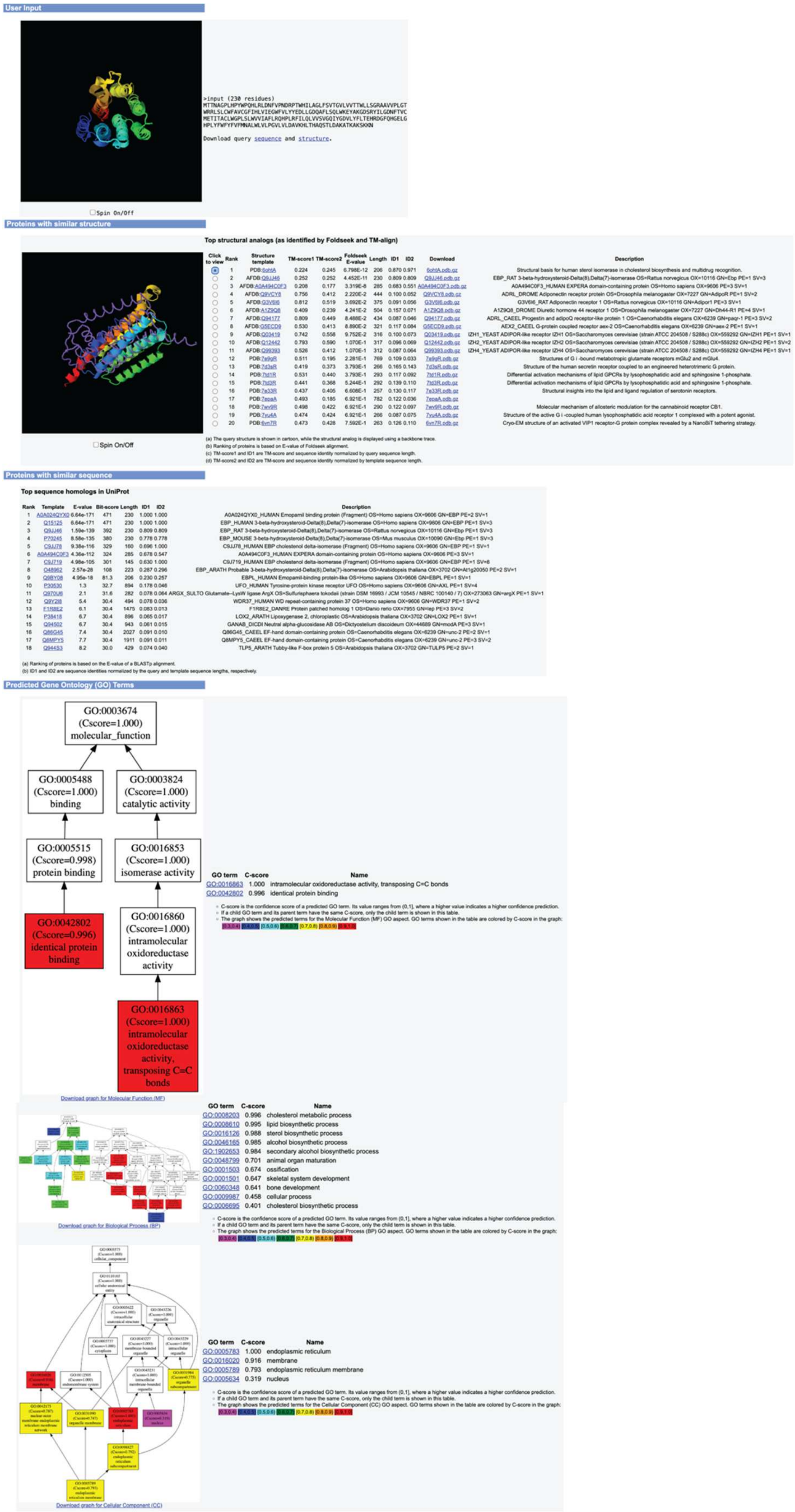


**Figure S8.** The details for each target protein in structure-function database.
